# The fitness consequences of wildlife conservation translocations: a meta-analysis

**DOI:** 10.1101/2023.01.14.524021

**Authors:** Iwo P. Gross, Alan E. Wilson, Matthew E. Wolak

## Abstract

Conservation translocation is a common strategy to offset mounting rates of population declines through the transfer of captive-or wild-origin organisms into areas where conspecific populations are imperiled or completely extirpated. Translocations that supplement existing populations are referred to as reinforcements, and can be conducted using captive-origin animals (*ex situ* reinforcements [ESR]) or wild-origin animals without any captive ancestry (*in situ* reinforcement [ISR]). These programs have been criticized for low success rates and husbandry practices that produce individuals with genetic and performance deficits, but the post-release performance of captive-origin or wild-origin translocated groups has not been systematically reviewed to quantify success relative to wild-resident control groups. To assess the disparity in post-release performance of translocated organisms relative to wild-resident conspecifics and examine the association of performance disparity with organismal and methodological factors across studies, we conducted a systematic review and meta-analysis of 821 performance comparisons from 171 studies representing nine animal classes (101 species). We found that translocated organisms have 64% decreased odds of out-performing their wild-resident counterparts, supporting claims of systemic issues hampering conservation translocations. To help identify translocation practices that could maximize program success in the future, we further quantified the impact of broad organismal and methodological factors on the disparity between translocated and wild-resident conspecific performance. Pre-release animal enrichment significantly reduced performance disparities, whereas our results suggest no overall effects of taxonomic group, sex, captive generation time, or the type of fitness surrogate measured. This work is the most comprehensive systematic review to date of animal conservation translocations in which wild conspecifics were used as comparators, thereby facilitating an evaluation of the overall impact of this conservation strategy and identifying specific actions to increase success. Our review highlights the need for conservation managers to include both sympatric and allopatric wild-reference groups to ensure the post-release performance of translocated animals can be evaluated. Further, our analyses identify pre-release animal enrichment as a particular strategy for improving the outcomes of animal conservation translocations, and demonstrate how meta-analysis can be used to identify implementation choices that maximize translocated animal contributions to recipient population growth and viability.

## I. INTRODUCTION

Rates of floral and faunal species loss in the new Anthropocene signal the onset of a sixth global mass extinction (Barnosky *et al*., 2011; Pimm *et al*., 2014). Among vertebrates, extinction rates in the last century increased 100 times above pre-industrial levels (Ceballos *et al*., 2015) and recorded population size declines have averaged 68% in the last 50 years (WWF/ZSL, 2020). Continued species loss will have irreversible consequences for ecosystem services and resiliency. Slowing the rate of species loss will require eliminating existing threats (e.g., disease, poaching;Heppell, Crowder & Crouse, 1996; Snyder *et al*., 1996) and extensive habitat preservation and restoration (Bouzat *et al*., 2009; Newmark *et al*., 2017). While environmental threats are being addressed, supportive breeding and translocation programs have been proposed to maintain species and their roles in the ecosystem (Ford, 2002; Seddon *et al*., 2014; Hufbauer *et al*., 2015).

Animal conservation translocations (ACTs) are a common strategy intended to directly offset species extinction or provide beneficial ecosystem functions through the supplementation of threatened populations and the repatriation of extirpated populations (Ceballos, Ehrlich & Dirzo, 2017; Ceballos, Ehrlich & Raven, 2020). ACT programs supplementing extant populations in their indigenous range are called reinforcements, and can be grouped into two categories (Seddon, Armstrong & Maloney, 2007; IUCN/SCC, 2013). First, *in situ* reinforcements (ISRs) collect and transport wild organisms to another natural area for the benefit of the recipient conspecific population. Second, *ex situ* reinforcements (ESRs) incorporate a captive phase lasting from less than one generation (i.e., head-starting) up to several generations (i.e., captive breeding) before release. Despite immense efforts across a wide array of taxa to employ ACTs and document their efficacy in the literature, a major unresolved question is whether translocated individuals are as capable as their wild conspecifics at contributing to population viability and preventing extinctions.

To date, ACTs have had many celebrated results and a quantifiable impact on global extinction rates (Hoffmann *et al*., 2010; Jachowski *et al*., 2011; Hess *et al*., 2012; Jensen *et al*., 2018; Bolam *et al*., 2021). Nevertheless, ACTs are often considered a halfway technology with logistical and systemic shortcomings that result in failure to achieve pre-defined conservation goals (Snyder *et al*., 1996; Fischer & Lindenmayer, 2000; Pérez *et al*., 2012; Germano *et al*., 2015; Sullivan, Nowak & Kwiatkowski, 2015). In particular, multiple studies indicate that ACTs might produce individuals with inherent genetic and performance-related deficits that compromise ACT program goals and reduce recipient population viability (Frankham, 2008; Satake & Araki, 2012; Farquharson, Hogg & Grueber, 2018, 2021). These deficits are most severe in ESRs involving multiple captive generations since the capture of ESR founders acts as a bottleneck on genetic diversity. Even with a stringent captive breeding structure, factors inherent to husbandry practices (e.g., small effective population size, inbreeding depression, relaxed purifying selection) increase the likelihood of fixation of undesirable alleles and a loss of allelic heterozygosity (Lynch & O’Hely, 2001; Ford, 2002; Araki *et al*., 2008; Furlan *et al*., 2020). A large influx of captive-born individuals into the wild with such genetic properties can swamp genetic diversity of locally adapted populations and reduce effective population sizes despite an overall increase in numbers (Ryman & Laikre, 1991; Alleaume-Benharira, Pen & Ronce, 2006; Laikre *et al*., 2010; Ford, Murdoch & Howard, 2012; Waples *et al*., 2016; Pinter, Epifanio & Unfer, 2019). For example, Klütsch *et al*., (2019) examined how stocking of captive-bred brown trout (*Salmo trutta*) influenced the genetic diversity of wild sub-populations isolated by hydroelectric dams. They found that historically stocked sections of the system exhibited genetic signs of bottleneck events indicative of genetic swamping by the released captive-bred fish. However, such genetic deficits require multiple generations of captive breeding to accumulate which suggests alternative factors contribute to fitness reductions observed in ESRs lasting a generation or less (Araki, Cooper & Blouin, 2007; Farquharson *et al*., 2018).

A fundamental criticism of ESRs is that non-natural conditions present in captivity can influence individual phenotypes, diminishing post-release performance relative to wild conspecifics (Araki *et al*., 2008; Stuparyk *et al*., 2018; Tetzlaff, Sperry & DeGregorio, 2019a; Crates, Stojanovic & Heinsohn, 2022), where animal performance is defined as a measure of some critical life history character (e.g., growth rate, survival, fecundity) that correlates with individual fitness (Thoday, 1953; Hendry *et al*., 2018). In a variety of taxa, ESRs lasting a generation or less negatively influence post-release performance metrics such as anti-predator behavior (Kraaijeveld-Smit *et al*., 2006; Melstrom, Salau & Shanafelt, 2019), dispersal (Lehrer *et al*., 2016; Degregorio *et al*., 2017; McCallen *et al*., 2018), growth (Horreo *et al*., 2018), survival (McCleery *et al*., 2013; Vanderwerf *et al*., 2014), and reproductive success (Araki *et al*., 2007; Knudsen *et al*., 2008; Ford *et al*., 2016). Such phenotypic mismatches can arise *via* domestication selection, where trait values that are selected for in captivity are maladaptive when expressed in a natural environment (Heath *et al*., 2003; Stringwell *et al*., 2014).

The unintentional selection for domesticated phenotypes in artificial environments is likely a major contributor to reduced performance of ESRs, since it can alter phenotype distributions within a single generation (Araki *et al*., 2007; Horreo *et al*., 2018). For example, Araki *et al*., (2007) discovered that captive-bred steelhead trout (*Oncorhynchus mykiss*) exhibited a 40% reduction in fecundity relative to their wild counterparts for every additional generation of captive ancestry. In the absence of domestication selection, phenotypic plasticity during less than one generation of captive development and rearing can lead to similar deviations from the wild-type in anatomical, physiological, and behavioral traits (Huntingford, 2004; Stamps & Swaisgood, 2007; Stringwell *et al*., 2014). Moreover, certain plastic phenotypic deviations have an epigenetic basis, thereby potentially influencing performance of future generations regardless of their developmental environment (Day & Bonduriansky, 2011; Evans *et al*., 2014).

By virtue of the use of wild-born organisms and lack of a captive component, ISRs are lauded as a less intrusive and a generally more successful translocation strategy than ESRs (Griffith *et al*., 1989; Wolf *et al*., 1996; Fischer & Lindenmayer, 2000; Kingsbury & Attum, 2009; Rummel *et al*., 2016). However, failure rates for ISRs are still notably high and potentially a consequence of similar issues that affect ESRs. At the population-level, supplementing existing populations with conspecifics from other wild populations can lead to maladaptive gene flow or outbreeding depression of the recipient population and a reduction in meta-population genetic diversity (Lenormand, 2002; Moritz, 2002; Ficetola & De Bernardi, 2005). At the individual-level, ISRs can impact post-release performance either directly through transportation-related stress (Dickens, Delehanty & Michael Romero, 2010) or indirectly through energetic costs of establishment (Armstrong & Seddon, 2008), errant dispersal and homing behaviors (Germano & Bishop, 2009), diminished social structure (Goldenberg *et al*., 2019), and phenotypic mismatches within the novel ecosystem (Stamps & Swaisgood, 2007; Turlure *et al*., 2013).

Exposing captive organisms to some form of enrichment is the most intuitive strategy to counteract phenotypic mismatch, and a reasonable compromise when ISRs are not possible. We use the term enrichment broadly to refer to a variety of supportive measures (Fischer & Lindenmayer, 2000) employed by ACTs to offset any maladaptive behavior, physiology, or physical injury incurred during captivity and/or translocation (Reading, Miller & Shepherdson, 2013; Goldenberg *et al*., 2019; Tetzlaff *et al*., 2019a). Some examples include fostering pre-release organisms with experienced conspecifics (Lumsden & Drever, 2002; Shier & Owings, 2007), simulating *in situ* environments with naturalistic enclosures or antipredator training (Brown, Davidson & Laland, 2003; Nazar & Marin, 2011; Roe, Frank & Kingsbury, 2015; Tetzlaff, Sperry & DeGregorio, 2018; Zhu *et al*., 2022), implementing a soft-release phase where supplemental feeding and/or shelter are provided (Tuberville *et al*., 2005; Brown *et al*., 2006), or simply reducing the length of the captive phase (Degregorio et al., 2017; DeGregorio, Weatherhead, Tuberville, & Sperry, 2013; but see Tetzlaff, Sperry, Kingsbury, & DeGregorio, 2019). Literature reviews suggest that enriched ESR cohorts are more successful post-release compared to un-enriched cohorts (Fischer & Lindenmayer, 2000; Tetzlaff *et al*., 2019a), but the lack of a wild-reference group in the synthesized findings makes it difficult to conclude whether enrichment strategies produce translocated organisms that are indistinguishable from wild conspecifics with respect to post-release fitness (Mathews *et al*., 2005).

Finally, the metric used to gauge relative performance can vary in its representation of fitness, thereby shaping study conclusions (Moyes *et al*., 2009; Wilson & Nussey, 2010; Hendry *et al*., 2018). The lifetime reproductive success of an individual is the ideal measure of fitness when focusing on population dynamics and phenotypic adaptation, but lifetime reproductive success and the component life history traits that determine it (e.g., survival and fecundity) are difficult to measure in natural conditions (Coulson *et al*., 2006; Fairbairn & Reeve, 2001; Hendry *et al*., 2018; Kozłowski, 1993). Consequently, fitness is often approximated using traits that typically correlate with fitness (e.g., body size, body constitution, movement). However, the extent to which correlates and components of fitness (hereafter collectively referred to as fitness surrogates) covary and ultimately relate to lifetime reproductive success remains uncertain (Moyes *et al*., 2009; Hendry *et al*., 2018). For example, Mulder *et al*., (2017) found that while translocated and wild-resident desert tortoises (*Gopherus agassizii*) had similar body condition and survivorship four years post-release, translocated males fathered none of 92 offspring for which paternity was determined. This suggests that survival is a potentially inappropriate component of fitness to gauge ACT success in adult organisms with high adult survivorship. Conservation translocations should consider multiple relevant fitness components during project assessment to identify potential life-history trade-offs and avoid misleading conclusions (Fincke & Hadrys, 2001; Wilson & Nussey, 2010; Pekkala, Kotiaho & Puurtinen, 2011; Hendry *et al*., 2018).

The translocation literature has grown to a point that quantitative reviews are necessary to summarize the high volume of papers, while also accounting for heterogeneity and bias (Bajomi *et al*., 2010; Stewart, 2010). Seminal reviews have evaluated ACT efficacy from the published literature and *via* mail-in questionnaires sent to conservation program managers (e.g., Griffith *et al*., 1989; Wolf *et al*., 1996b; Fischer & Lindenmayer, 2000; Kraaijeveld-Smit *et al*., 2006; Seddon *et al*., 2007; Rummel *et al*., 2016; Brichieri-Colombi *et al*., 2019). In these approaches, the response variable (translocation success *versus* failure) reduces a highly nuanced outcome to a binary classification based in part on unstandardized criteria (e.g., creation of a self-sustaining population) and professional opinion. These syntheses are an invaluable resource for applied conservation and management (Nason *et al*., 2021), but their largely narrative format is not conducive to quantitative synthesis across broad and multifactorial datasets while accounting for bias and heterogeneity among studies.

Meta-analysis is a quantitative tool that integrates the findings of published papers to examine broader patterns and draw more powerful conclusions about a focal topic (Glass, 1976; Stewart, 2010; Gurevitch *et al*., 2018). Meta-analyses standardize and pool effect sizes from multiple studies while preserving the magnitude and variance of the original effects, thus enabling predictions about particular driving variables to be modeled as covariates that account for unexplained variation among studies (Glass, 1976; Bennett *et al*., 2017). A properly executed meta-analysis of the conservation literature will synthesize existing trends, assess knowledge gaps, and develop recommendations for future research and management actions.

Meta-analysis is well-equipped to gauge translocation efficacy for establishing viable populations based on the post-release performance of animal subjects (Ducatez & Shine, 2019; Jule, Leaver, & Lea, 2008; Linklater *et al*., 2011; Skikne *et al*., 2020; Tetzlaff, Sperry, & DeGregorio, 2019; Zhang, Gao, & Zhang, 2022; Zhu *et al*., 2022; also see Resende *et al*., 2021). Previous meta-analyses fail to include a wild-reference group (but see Stuparyk *et al*., 2018) and therefore cannot address the way by which translocated individuals may contribute to the population viability goals of ACTs. For instance, translocated animals that are on average less likely to survive, reproduce, or perform relative to wild conspecifics in the recipient population (i.e., a sympatric control group comparison) will have limited contributions to genetic or evolutionary mechanisms increasing population persistence (i.e., genetic and evolutionary rescue; Carlson *et al*., 2014). The performance of wild members of the recipient population should be considered by conservation managers as an adaptive baseline shaped by local environmental and social pressures that translocation programs should strive to emulate (Mathews *et al*., 2005). Even though translocated cohorts might exhibit lower survival or reproductive success compared to wild control groups, translocations may positively affect other demographic and ecological processes (e.g., demographic rescue and Allee effects) that fulfill primary goals of ACTs (IUCN/SSC, 2013). Distinguishing these beneficial effects on the wild resident recipient population requires comparison with wild allopatric control groups to separate the effect of translocation from the measurement of performance in both the translocated cohort as well as the wild sympatric recipient group. However, use of both wild allopatric and sympatric control groups has rarely been reported (e.g., Molony *et al*., 2006; Sah *et al*., 2016).

Here, we present the first comprehensive meta-analysis to assess conservation translocation programs through the synthesis of studies involving direct, post-release comparisons of translocated and wild-resident conspecifics. We further expand on previous reviews by sampling the literature broadly across macroscopic animal groups and assessing relative post-release performance using multiple fitness surrogates. Our primary objective is to quantify the disparity in post-release performance of translocated organisms relative to wild-resident conspecifics across animal taxa and ultimately gauge the potential contribution of translocated individuals to genetic or evolutionary rescue strategies. We performed a systematic review and meta-analysis to quantify this performance disparity and also sought to determine whether organismal and methodological factors influence this disparity by extracting information about (*i*) sex and (*ii*) taxonomy as well as (*iii*) what strategies were implemented, if at all, regarding pre-release animal enrichment and categorizing (*iv*) translocation reinforcements as *in situ* vs. *ex situ*, (*v*) captive generation time, and (*vi*) the type of fitness surrogate measured. Our systematic review and meta-analysis were guided by how we predict the above factors will influence the relative performance among translocated and wild cohorts:

1. Animals incorporated into conservation translocation programs represent a wide taxonomic breadth in which natural history, sensory capabilities, and native environments will vary substantially. These factors will also affect the capacities of conservation managers to recreate natural conditions in captivity and in turn influence organismal response. We predict that taxonomic groups will broadly vary in their relative post-release performance.
2. Disparity in performance among translocated and wild-resident organisms will increase as a function of whether a trait is considered an indirect correlate (e.g., movement and body constitution) or a direct component of an organism’s fitness (e.g., growth, survival, reproductive success).
3. Within species, sexes can vary in physical form and physiology, seasonal and annual energetic investments, optimal life-history strategy, spatial ecology, and behavior (Trivers, 1972; Parker, 1979; Ruckstuhl & Neuhaus, 2005; Bonduriansky & Chenoweth, 2009; Fairbairn, 2013). We predict significant intraspecific variation in post-release relative performance among male and female cohorts.
4. We predict varying amounts of captive ancestry will explain variation in relative post-release performance of translocated cohorts. We predict that ESRs lasting > 1 generation will sustain greater disparity in relative performance than ESRs lasting < 1 generation (e.g., head-starting programs). We also predict that ISRs involving zero captive generations will have more similar post-release performances to wild-reference groups compared to ESRs.
5. Translocated organisms exposed to one or more forms of enrichment will perform better post-release relative to wild conspecifics compared to unenriched translocated conspecifics.

## II. METHODS

### (1) Literature search and inclusion criteria

We conducted our literature review, screening, and analysis according to Preferred Reporting Items for Systematic Reviews and Meta-Analyses (PRISMA) guidelines (Fig. 1; Moher *et al*., 2009) and following a PRISMA EcoEvo checklist (Appendix S1; O’Dea *et al*., 2021). We searched the literature using Web of Science, including all databases (01/01/1900-08/23/2019), for peer-reviewed studies concerning animal conservation translocations with some reference to a wild-resident group used as a control. Our query was:

**Figure 1.**
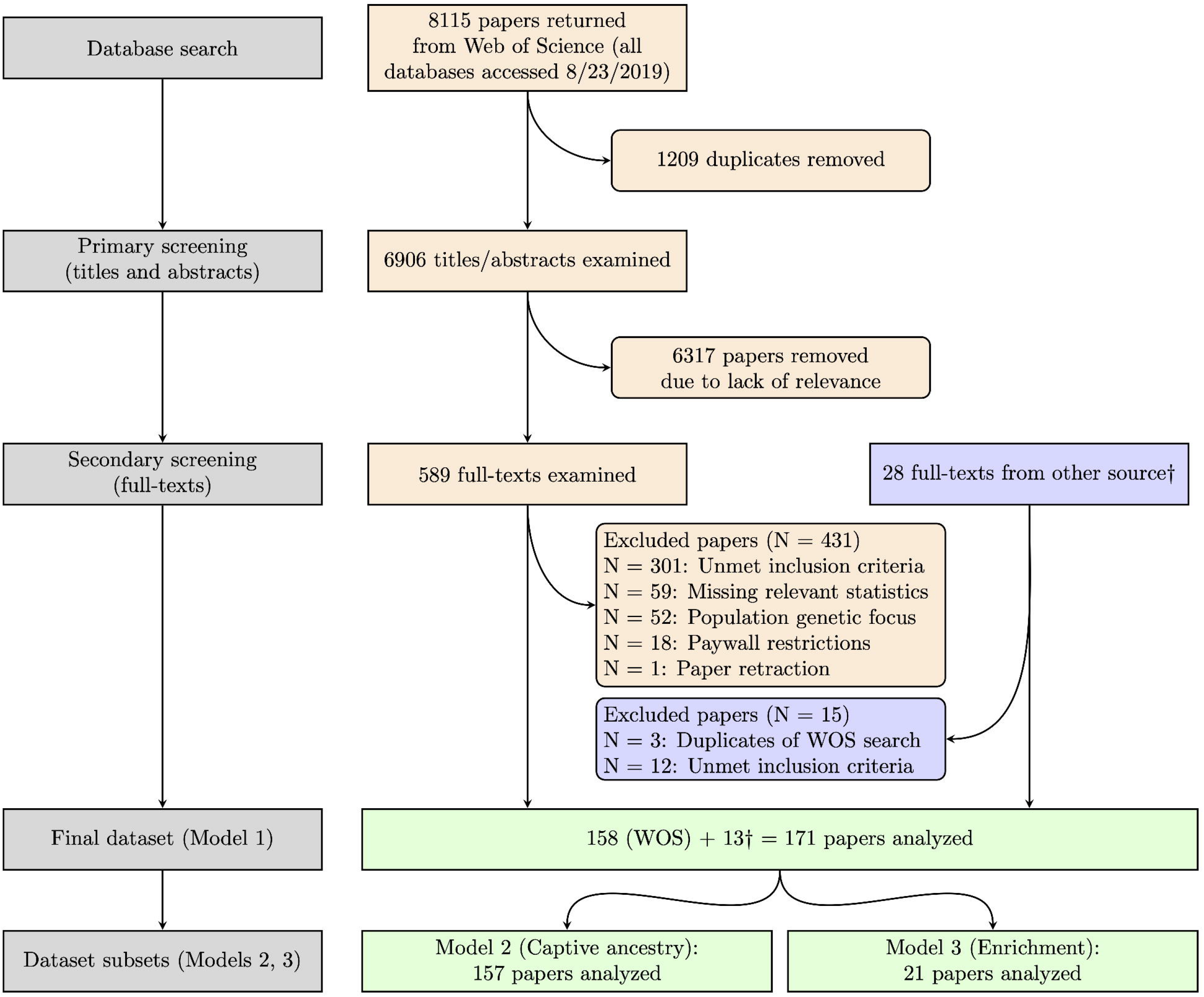
A flowchart describing the dataset accession and screening processes as prescribed by Preferred Reporting Items for Systematic Reviews and Meta-Analyses (PRISMA) guidelines (Moher *et al*., 2009; O’Dea *et al*., 2021). †We supplemented the query-returned papers with 28 additional papers synthesized by Stuparyk *et al*., (2018) focused on relative post-release performance of organisms translocated to alleviate human-animal conflicts.

TS = ((conservation OR reintroduc* OR repatriat* OR releas* OR supplementation OR transloc*) AND (“captive born“ OR “captive breed*“ OR “captive bred“ OR “captive rais*“ OR “captive rear*“ OR fisher* OR hatcher* OR “headstart*“ OR “head start*“ OR relocat*) AND (native OR resident OR wild) AND (animal OR invertebrate OR fish* OR amphib* OR reptil* OR mammal* OR bird* OR avian*))

We also screened studies reviewed in Stuparyk *et al*., (2018) for inclusion because our primary Web of Science query did not encompass the terminology used in several ISR studies concerning ISRs intended to mitigate human-animal conflicts. After screening (Fig. 1; Table S1 in Appendix S2), the resulting 589 full-text papers—along with 28 additional papers synthesized by Stuparyk *et al*., (2018)—were assessed for our required inclusion criteria: At least one direct, quantitative comparison of a fitness component (growth, survival, reproduction) or fitness-correlated trait (movement, body constitution) between one translocated (either ESR or ISR) and one wild-resident animal group. The comparison(s) must have been made in a natural setting following the release of the translocated cohort into an area occupied by wild-resident conspecifics of comparable age and size distributions. We excluded papers making comparisons based on ambiguous traits without the author(s) clearly defining a relationship to fitness. These ambiguous traits include, but are not limited to, stable isotope or itemized diet analysis (e.g., Bourass & Hingrat, 2015; Quinn, Seamons, & Johnson, 2012), habitat use (e.g., Himes, Hardy, Rudolph, & Burgdorp, 2006; Mann, Marty Holtgren, Auer, & Ogren, 2011), behavioral ethograms (e.g., Rantanen, Buner, Riordan, Sotherton, & Macdonald, 2010), or varying morphotypes (Larsen *et al*., 2013). Papers were also excluded if they did not contain the necessary descriptive statistics to calculate effect sizes (see “effect size extraction” below) and corresponding authors did not respond to email requests for supplementary statistics. Finally, we excluded papers assessing the influence of translocations on population genetic indices, as these datasets do not involve distinct animal groups and sample sizes that are necessary to calculate effect sizes.

### (2) Effect size extraction and calculation

An effect size is a statistic used to estimate the magnitude of a relationship between two experimental groups or variables (Nakagawa & Cuthill, 2007). We quantified the disparity in performance between wild-resident and translocated cohorts using two effect size metrics. In studies comparing animal groups based on mean estimates of continuous traits (*e.g.,* growth rate, survivorship, clutch size), we calculated standardized mean differences (SMD; Rosenthal & Rubin, 1982). For comparisons based on counts of dichotomous outcomes (e.g., full vs. empty stomachs, proportion recaptured, successful vs. unsuccessful nests), we calculated log odds ratios (LOR; Haddock, Rindskopf, & Shadish, 1998). Effect sizes were calculated using the function *escalc* in the R package *metafor* (Viechtbauer, 2010). From each paper, we extracted descriptive statistics necessary to calculate effect sizes for all relevant fitness-related comparisons from the published and supplemental texts and data tables, from additional materials requested from corresponding authors, and from relevant figures and plots using the R package *digitize* (Poisot, 2011).

We converted all SMD estimates to LORs for more intuitive interpretation of effect size magnitudes (Chinn, 2000; Rosenberg, Rothstein & Gurevitch, 2013). Log odds ratios are centered at zero and were calculated such that positive values indicate translocated individuals exhibited better relative performance than wild-resident individuals, and negative values indicate translocated individuals under-performed relative to wild individuals. In traits where lower values indicate better performance (e.g., rate of mortality, parasite load), raw trait values were multiplied by negative one for consistent interpretation.

### (3) Moderators

We extracted several study-level and estimate-level attributes to evaluate our predictions in the final statistical model. Comparisons were assigned a study ID based on the paper from which they were extracted and a group ID to distinguish unique groups of organisms within each study. Identical group ID values were assigned to comparisons involving measures of different traits in the same organisms (Noble *et al*., 2017). We also retrieved taxonomic class, order, family, genus, and species names of all assessed organisms from the Integrative Taxonomic Information System database (ITIS, 2021). We identified the fitness-related trait under consideration in each comparison and reclassified each into one of five broad categories: Movement, body constitution, growth, survival, and reproduction. We also identified the predominant sex (i.e., female-only, male-only, or mixed-sex) of organisms in each comparison. We characterized the amount of captive ancestry of translocated individuals represented in each comparison using two separate variables. First, we distinguished comparisons according to whether the translocation program involved a captive phase or not (i.e., *in situ* reinforcement vs. *ex situ* reinforcement strategy). In those studies that described the amount of captive ancestry among translocated cohorts, we further classified captive ancestry using an ordered set of categories: 1) ISR organisms without any captive ancestry, 2) ESR organisms with wild-origin parents not raised in captivity, and 3) ESR organisms with one or more direct ancestral generation raised in captivity. Finally, we classified the absence/presence (0/1) of enrichment techniques for select comparisons according to the circumstances of each paper. Due to the arbitrary nature of enrichment treatments across our dataset, we only assessed enrichment efficacy within studies that included post-release performance comparisons with wild-resident conspecifics for both enriched and unenriched translocated cohorts.

### (4) Statistical analysis

#### (a) Model structure

We first fit a meta-analytical taxonomic mixed model (Clutton-Brock & Harvey, 1977; Hadfield & Nakagawa, 2010) to quantify the overall effect size—relative post-release performance among translocated and wild-resident cohorts—across the entire dataset, and also assess how individual-level (i.e., sex, taxonomic group) and study-level (i.e., fitness surrogate, ESR vs. ISR strategy) factors influence this outcome (model 1; Table S2 in Appendix S2). We modeled the LOR responses as a function of categorical fixed effects for sex, taxonomic class, fitness surrogate, and study strategy. We examined two-way interactions between sex and fitness surrogate, and between sex and strategy to determine whether these individual- and study-level factors had sex-specific relationships with relative post-release performance. We also included a two-way interaction between fitness surrogate and strategy to examine whether post-release performance in various fitness-related traits is influenced by translocation strategy.

#### (b) Quantification of study heterogeneity

We included several random effects terms to account for different sources of non-independence among effect sizes. To account for shared taxonomy (Clutton-Brock & Harvey, 1977; Hadfield & Nakagawa, 2010), we included all Linnaean taxonomic ranks below the level of class as separate random effect terms in each model. Taxonomic class was maintained as a fixed effect to quantify average differences in post-release performance across broad, recognizable animal groups of potential relevance to translocation managers. We chose the nested taxonomic random effects approach over the incorporation of a phylogenetic covariance matrix because the taxonomic breadth of our dataset precluded access to a comprehensive phylogeny with empirically-derived branch lengths (Hadfield & Nakagawa, 2010; Lajeunesse, Rosenberg & Jennions, 2013) and the goals of our analysis are not concerned with estimating the extent of correlated evolution. We included study and group identities as two additional random effects terms to account for non-independence of effect sizes within studies and among repeated measures of unique organisms (Noble *et al*., 2017). We also included a term to model the variance in sampling errors for each effect size, which incorporates study-level precision into the model (see “*Meta-analysis model specifications*” below; Hadfield, 2010; Mengersen *et al*., 2013). To assess the overall dispersion of our effect sizes and thus the generalizability of the findings of our models, we calculated total model heterogeneity (Higgins & Thompson, 2002; Table S5 in Appendix S2). Since measures of heterogeneity tend to be high in ecological studies assessing diverse taxa (Senior *et al*., 2016), we further calculated the heterogeneity apportioned across the aforementioned taxonomic and study-specific random effect terms to more clearly explain patterns of dispersion in effect sizes (Nakagawa & Santos, 2012).

#### (c) Meta-analysis model specifications

The model was fit in a Bayesian framework using a Markov chain Monte Carlo (MCMC) algorithm as implemented in the R package *MCMCglmm* (Hadfield, 2010). We specified diffuse normal prior distributions for all fixed effects (mean=0, variance=10^10^) and parameter expanded priors on all variance components to yield scaled non-central F-distributions with numerator and denominator degrees of freedom equal to one and a scale parameter of 1000 (Gelman, 2006; Hadfield, 2010). The model was constructed as a random-effect meta-analysis by including a term that contained the estimated sampling variance for each effect size and fixing the variance associated with this term to one (in the prior). The MCMC chain for each model was run for an initial burn-in of 3000 iterations with samples from the chain being saved every 150^th^ iteration until 3000 samples were obtained. All models were checked for autocorrelation values < |0.1|, and model convergence was confirmed using a Heidelberg stationarity test and through visual assessment of posterior distributions.

#### (d) Influence of captive ancestry and enrichment

To assess the potential influence of domestication selection on the relative post-release performance of translocated cohorts, we used a second, similar model to analyze a subset of studies from the model 1 dataset that specified the number of captive generations experienced by translocated cohorts (model 2; Table S3 in Appendix S2). We fit model 2 by substituting the fixed effect of ESR vs. ISR strategy from model 1 with captive ancestry, a categorical fixed effect with three levels classifying the focal ACT cohort as either part of an ISR (without any captive ancestry), a head-starting program (ESR with < 1 generation in captivity), or a captive-breeding program (ESR with > 1 direct ancestral generations in captivity). We were unable to analyze captive ancestry as a continuous variable due to a lack of reporting of exact ancestral generation times, and due to cohorts of potentially mixed ancestry. Model 2 also included categorical fixed effects for sex, taxonomic class, fitness surrogate, and captive ancestry, and two-way interactions between sex and fitness surrogate, sex and captive ancestry, and between fitness surrogate and captive ancestry.

Finally, we developed model 3 to test the influence of enrichment strategies on relative post-release performance among wild and translocated cohorts by analyzing a subset of studies from the model 1 dataset that gathered relative post-release performance data with a wild-control group from both enriched and unenriched cohorts within the same experimental design (Table S4 in Appendix S2). In model 3, we included categorical fixed effects for sex, taxonomic class, fitness surrogate, ESR vs. ISR strategy, and enrichment. We also included two-way interactions between enrichment and sex, enrichment and surrogate, enrichment and strategy.

#### (e) Results reporting

We addressed whether conservation programs produce organisms that perform similarly to or better than wild-resident conspecifics after release by calculating the total probability that the log odds ratio for a particular comparison is greater than or equal to zero. We also reported the marginal posterior mode and 95% highest posterior density credible interval (CrI) of the odds ratio (OR) by exponentiating posterior distributions of log odds ratios. We further interpreted each effect size according to established threshold values delimiting small, medium, and large magnitudes of effect (see Table 1; Cohen, 1992; Olivier *et al*., 2013). While these values provide potential useful benchmarks for the assessment of biological importance in effect sizes (Nakagawa & Cuthill, 2007), we focused our interpretations on the true effect size values and associated credible intervals. We used the R package *emmeans* to report the marginal values of individual parameters and overall model effects to account for the influence of other model parameters (Lenth, 2021). All derived parameters are calculated across the entire posterior distribution to carry through all uncertainty.

**Table 1.** Established recommendations for the interpretation of log odds ratio (LOR) and odds ratio (OR) effect size magnitude and direction (Cohen, 1992; Olivier *et al*., 2013). Effect size values in gray describe ACT comparisons where translocated cohorts have decreased odds of out-performing their wild-resident counterparts.

| Magnitude, direction of effect | Log odds ratio (LOR) | Odds ratio (OR) |
| --- | --- | --- |
| Large, negative | -1.10 | 0.33 |
| Medium, negative | -0.62 | 0.54 |
| Small, negative | -0.20 | 0.82 |
| No effect | 0 | 1.00 |
| Small, positive | 0.20 | 1.22 |
| Medium, positive | 0.62 | 1.86 |
| Large, positive | 1.10 | 3.00 |

### (5) Meta-analysis validation

#### (a) Outlier identification

We performed several meta-analysis validation tests to assess our dataset and findings for influential outliers and publication bias (Koricheva, Jennions & Lau, 2013). We identified outliers by calculating studentized deleted residuals for each comparison (Viechtbauer & Cheung, 2010). Comparisons with estimates over 1.96 standard deviations from the overall mean effect were considered outliers. We reran all models with outliers excluded to examine whether our findings were significantly affected.

#### (b) Publication bias

To test for temporal patterns in effect sizes and potential time-lag bias (Jennions & Møller, 2002; Koricheva *et al*., 2013), we fit a linear model with our dataset’s calculated effect sizes as the response variable and year of publication as the predictor. We included study ID, group ID, and all Linnaean taxonomic ranks below the level of phylum as random effects. To assess the dataset for evidence of publication bias, we used Egger’s regression to model the relationship of the meta-analytic residuals of each effect size against their precision (Egger *et al*., 1997; Nakagawa & Santos, 2012). Funnel plots were also created to visualize this relationship and identify any systematic absence of values, which could also indicate publication bias. To directly test the influence of publication bias on our overall effect, we conducted a trim-and-fill analysis using the *trimfill* function in the R package *metafor*, which quantifies funnel plot asymmetry and estimates the number of missing studies on either side of the overall mean effect (Duval & Tweedie, 2000; Viechtbauer, 2010).

## III. RESULTS

### (1) Description of data

Our initial Web of Science query returned 6906 unique articles. After filtering based on the inclusion criteria, our dataset comprised 821 unique effect sizes for relative performance of translocated organisms and their wild-resident conspecifics collected from 171 peer-reviewed studies (Table S1 in Appendix S2). The studies investigated 101 unique species in either *in situ* reinforcements (ISR; 599 effect sizes) or *ex situ* reinforcements (ESR; 222 effect sizes; Table 2). We calculated standardized effect sizes as log odds ratios (LORs) of performance metrics for translocated *versus* wild-resident cohorts such that positive values indicate translocated individuals exhibited higher relative performance than wild-resident conspecifics.

**Table 2.** Total number of log odds ratio (LOR) estimates collected, organized by taxonomic class and the fitness surrogate used as a metric for performance.

|  | Movement | Constitution | Growth | Survival | Reproduction | **Total** |
| --- | --- | --- | --- | --- | --- | --- |
| Bivalvia | 0 | 0 | 0 | 6 | 0 | **6** |
| Gastropoda | 0 | 3 | 5 | 9 | 6 | **23** |
| Malacostraca | 0 | 4 | 0 | 14 | 5 | **23** |
| Echinoidea | 1 | 1 | 0 | 0 | 0 | **2** |
| Actinopterygii | 38 | 43 | 26 | 158 | 77 | **342** |
| Amphibia | 4 | 2 | 0 | 3 | 0 | **9** |
| Reptilia | 101 | 12 | 26 | 54 | 9 | **202** |
| Aves | 14 | 12 | 0 | 59 | 38 | **123** |
| Mammalia | 18 | 13 | 2 | 46 | 12 | **91** |
| **Total** | **176** | **90** | **59** | **349** | **147** | **821** |

### (2) Meta-analysis

We analyzed the LORs and subsets of these data using three linear mixed effects meta-analytical models with fixed effect moderator variables to determine the impact of organismal and methodological factors plus random effects to attribute sources of variation among effect sizes.

#### (a) Model 1: Overall model

The primary model’s overall log odds ratio (LOR mode: -1.02; 95% CrI: -1.59 to -0.32) was converted to an odds ratio by exponentiating across the posterior distribution, indicating that translocated organisms have 63.9% decreased odds of out-performing their wild-resident counterparts (OR mode: 0.36; 95% CrI: 0.16 to 0.66). These effect size values correspond to a large magnitude of effect (Cohen, 1992; Olivier *et al*., 2013). By quantifying the proportion of posterior samples with LOR estimates greater than or equal to zero, we infer that translocated organisms had an overall 0.001 probability of out-performing wild-resident conspecifics following release (Fig. 2). Disparity in post-release performance did not vary widely across taxonomic classes, but instead all classes generally exhibited negative LOR effect sizes of moderate-to-high magnitude, suggesting poor performance of translocated organisms relative to wild-residents across taxa (Table S2 in Appendix S2). Similarly, relative post-release performance was largely unaffected by ESR vs. ISR translocation strategy.

**Figure 2.**
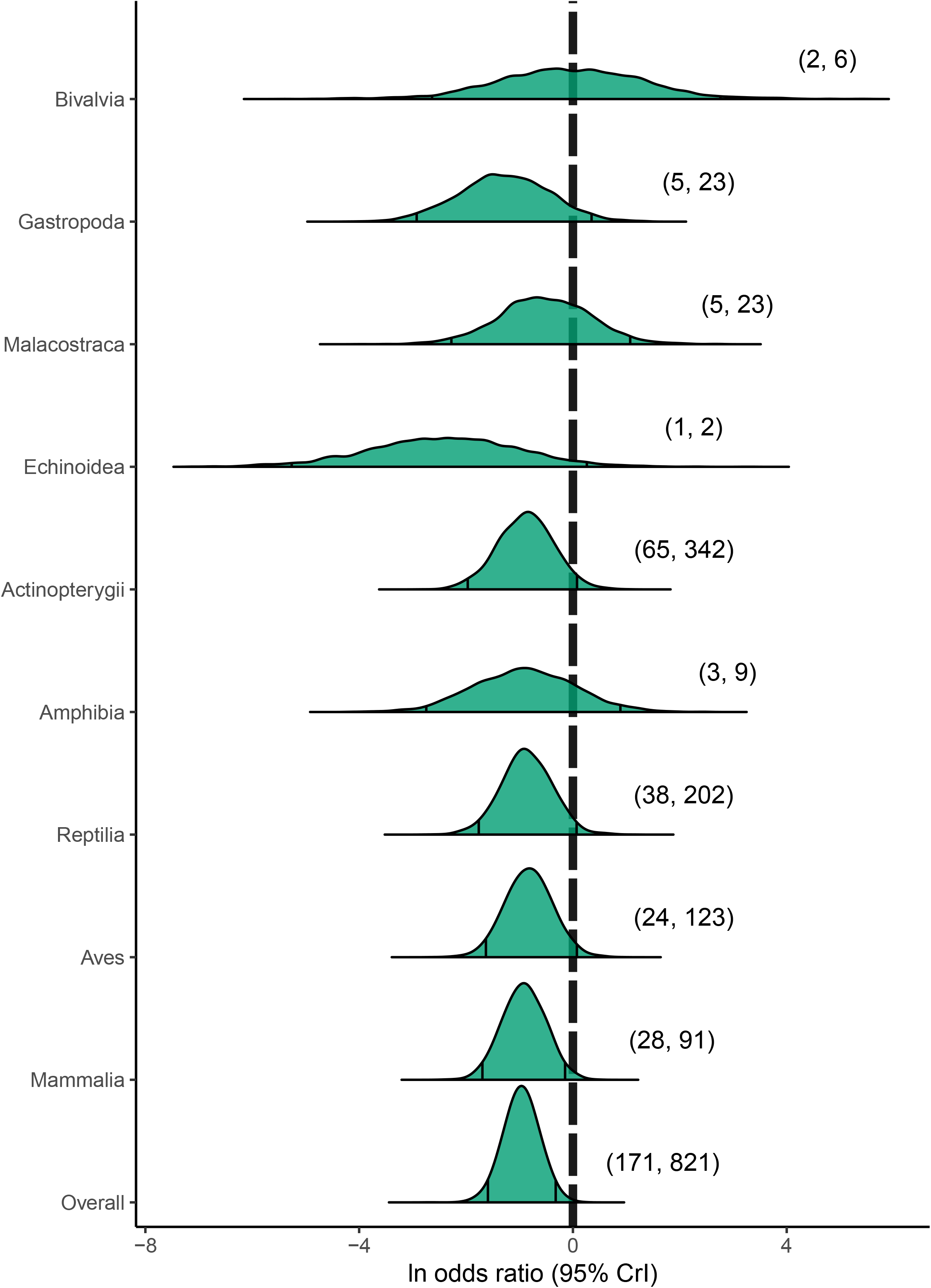
Marginal posterior distributions of log odds ratios (ln odds ratio) across taxonomic classes. Values greater than zero (vertical dashed line) indicate that translocated organisms out-performed wild-resident conspecifics, and negative values indicate translocated organisms under-performed. Vertical lines within each density plot indicate the 95% highest posterior density credible intervals (95% CrI). The parenthetical statements indicate the number of studies and the number of individual comparisons comprising each density plot, respectively.

In studies involving mixed-sex cohorts, comparisons based on fitness components tend to have lower LOR marginal posterior mode estimates than comparisons based on fitness correlates, but widely overlapping credible intervals across fitness components and correlates indicate a consensus in the interpretation of relative under-performance of translocated cohorts across a variety of performance metrics (Table S2 in Appendix S2). More variable effect size outcomes observed in sex-specific comparisons might suggest the presence of unequal life-history consequences to translocation among sexes, but low sample sizes in constitution-, growth-, and survival-based estimates, and uneven taxonomic representation in movement-based estimates preclude conclusive interpretations.

#### (b) Dataset heterogeneity

Here we summarize the heterogeneity statistics from model 1, representing the full dataset (see Table S5 in Appendix S2 for heterogeneity summary statistics of all three models). Heterogeneity values from individual taxonomic levels were relatively small and even in magnitude, but their combined taxonomic signal accounted for 60.6% (95% CrI: 47.0% to 74.7%) of total heterogeneity. Study- and group-level effects accounted for 23.9% (95% CrI: 12.0% to 34.3%) and 13.3% (95% CrI: 8.67% to 19.7%) of total heterogeneity, respectively.

#### (c) Subset analyses

Due to limited reporting of key moderator variables across studies, we analyzed subsets of the data in two additional models to assess whether the amount of captive ancestry or the presence of enrichment, respectively, influenced relative post-release performance of translocated cohorts. In general, LOR marginal posterior mode estimates were similar for the overall model effects and for parameters that also appeared in the primary model 1 above (Tables S3-S4 in Appendix S2). Therefore, we limit the remainder of our reporting to model-specific parameters.

##### (i) Model 2: Influence of captive ancestry

Post-release performance was largely unaffected by the length of captive ancestry, except for comparisons based on constitution and growth. In ISR studies involving constitution-based comparisons, we observed that relocated organisms without any captive ancestry had a lower probability (0.00; OR mode: 0.07; OR 95% CrI: 0.03 to 0.21) of out-performing wild-resident conspecifics relative to head-started (0.15; OR mode: 0.55; OR 95% CrI: 0.22 to 1.28) and captive-bred cohorts (0.25; OR mode: 0.21; OR 95% CrI: 0.02 to 2.51; Table S3 in Appendix S2). Comparisons with growth-based estimates exhibited a similar pattern, but these growth-related comparisons for relocated, head-started, and captive-bred cohorts were based on reduced datasets of only two, eight, and one studies, respectively.

##### (ii) Model 3: Influence of pre-release enrichment

To test the influence of enrichment strategies on relative post-release performance among translocated and wild cohorts, we analyzed a subset of studies that gathered data from both enriched and unenriched cohorts within the same experimental design and compared their post-release performances relative to a common wild-control group. Unenriched translocated organisms had a 0.06 probability of out-performing their wild-resident conspecifics (OR mode: 0.06; 95% CrI: 0.002 to 1.06), whereas enriched translocated organisms had a 0.58 probability of out-performing their wild counterparts (OR mode: 0.75; 95% CrI: 0.008 to 9.36; Fig. 3; Table S4 in Appendix S2).

**Figure 3.**
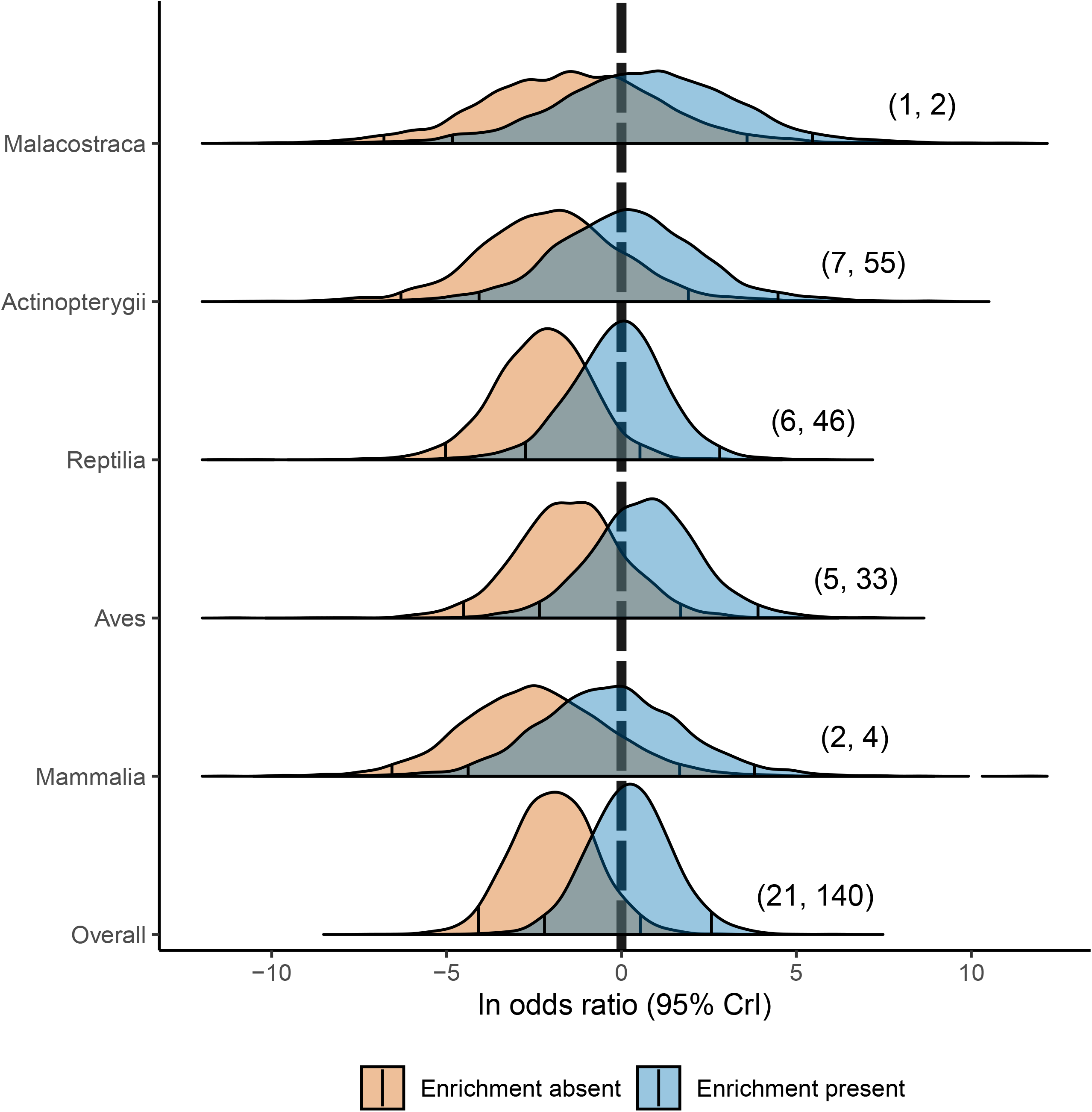
Marginal posterior distributions of log odds ratios (ln odds ratio) across enriched vs. unenriched translocated cohorts and taxonomic classes. Values greater than zero (vertical dashed line) indicate that translocated organisms out-performed wild-resident conspecifics, and negative values indicate translocated organisms under-performed. Blue density plots represent comparisons in which translocated organisms received some form of pre-release enrichment, whereas tan density plots represent comparisons where translocated organisms were unenriched. Vertical lines within each density plot indicate the 95% highest posterior density credible intervals (95% CrI). The parenthetical statements indicate the number of studies and the total number of comparisons between enriched/unenriched and wild cohorts organized by taxonomic class.

#### (d) Validation

##### (i) Outlier identification

We identified 56 outlying effect sizes based on studentized deleted residuals greater in magnitude than 1.96. Removal of outliers generally reduced effect sizes across all parameters and caused the 95% credible intervals for the overall effects of model 1 and model 2 to overlap with zero (Table S6-S8 in Appendix S2). However, outlier exclusion did not qualitatively affect our overall findings nor the indices of publication bias. Moreover, the examination of all outlying effect sizes revealed these effect sizes were calculated correctly and reflect legitimate patterns in relative post-release performance (Aguinis, Gottfredson & Joo, 2013). Therefore, we completed the final analysis with outliers included.

##### (ii) Publication bias

Visual examination of the marginal posterior distributions across years shows consistent effect sizes, indicating no time-lag bias in our dataset (Fig. 4). Removing outliers did not affect patterns of effect size magnitude across years (Fig. S1 in Appendix S3).

**Figure 4.**
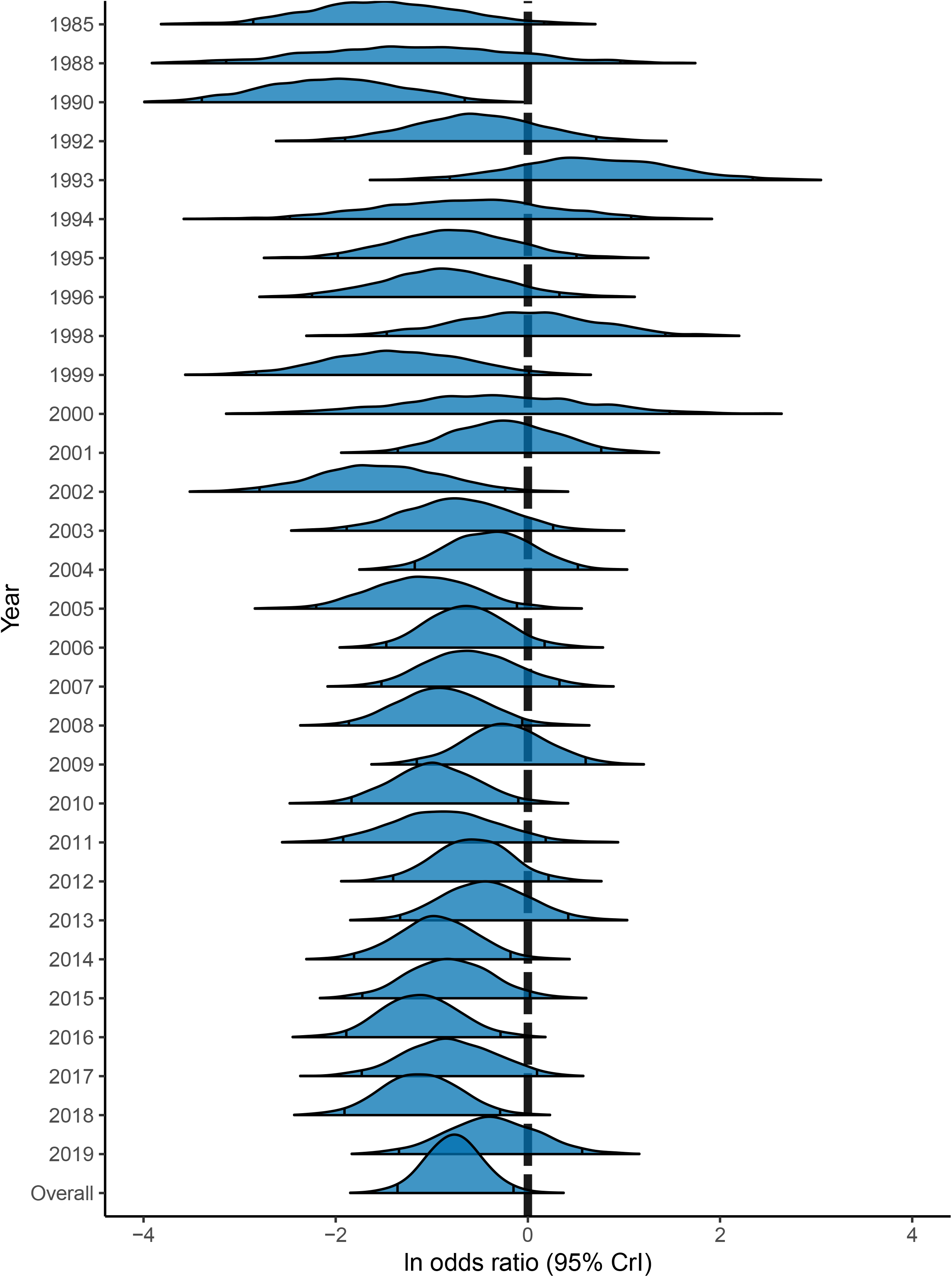
Time-lag bias. The relationship between effect size and publication year. Effect sizes are marginal posterior distributions, with 95% credible intervals indicated by black vertical lines. The dashed vertical line indicates an effect size of zero.

The Egger’s regression slopes were significantly different from zero after being run with both the dataset including (slope = 0.18; SE: 0.04; p < .001) and excluding (slope = 0.19; SE: 0.03; p < 0.001) outliers, which suggests the presence of publication bias. The resulting funnel plots revealed the presence of some effect size asymmetry in the dataset including outliers (Fig. 5), and to a lesser degree in the dataset excluding outliers (Fig. S2 in Appendix S3). However, the significant Egger’s regression slopes for both datasets could be an artifact of the high model heterogeneity and not publication bias (Higgins & Thompson, 2002).

**Figure 5.**
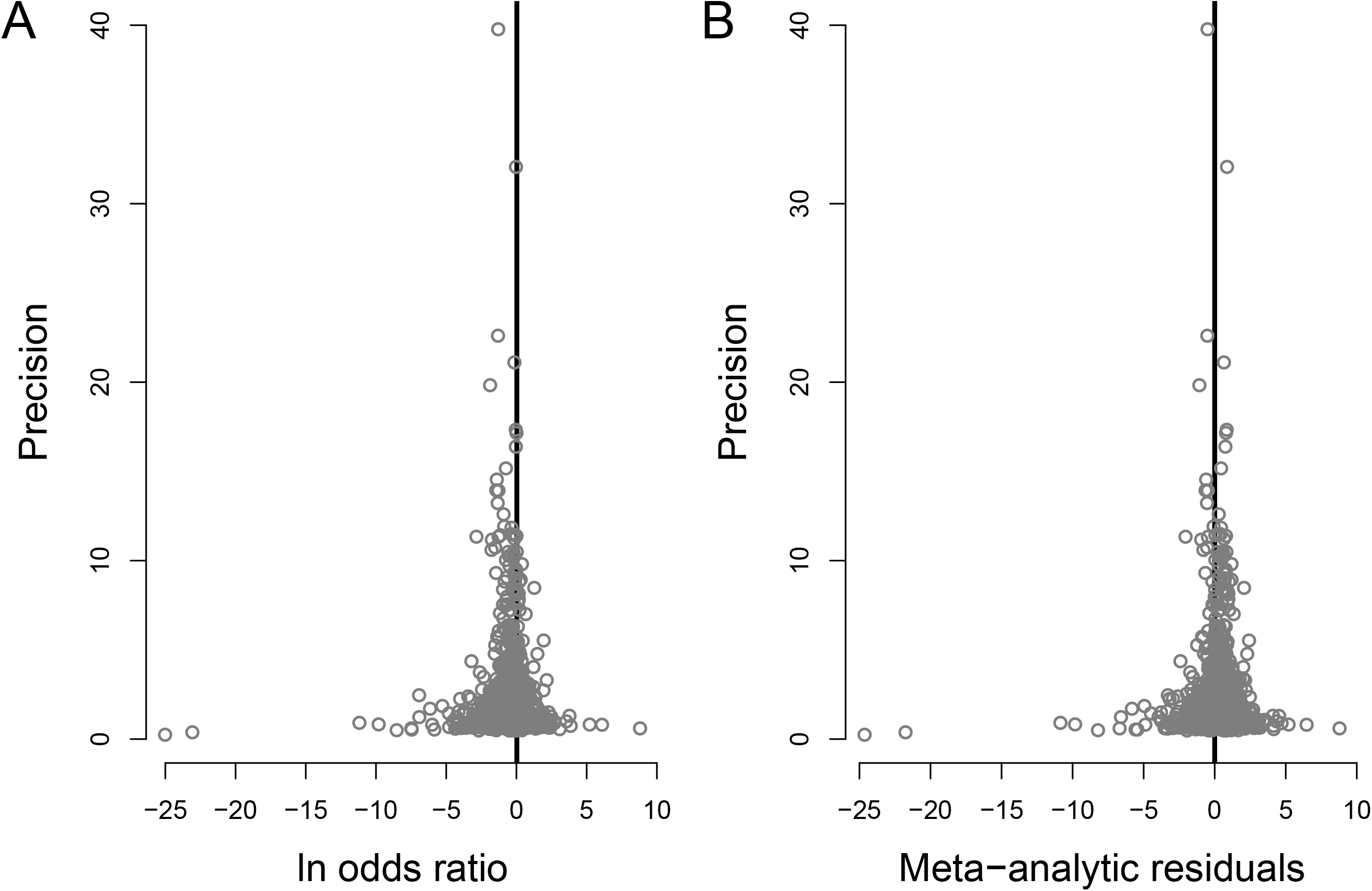
Egger’s regression. Funnel plots describing patterns of publication bias for the full dataset. (A) Relationship between effect size (log odds ratio) and precision. The solid line indicates the meta-analytic mean. (B) Relationship between the meta-analytic residuals and precision. The solid vertical line indicates zero.

The trim-and-fill analysis estimated that the overall model including outliers was missing zero estimates to the left side of the mean and 3 estimates from the right side (SE: 2.83; p = 0.06), whereas the model excluding outliers had zero missing estimates to either side of the mean (SE: 1.41; p = 0.5).

In conclusion, our validation tests indicate negligible amounts of publication bias. Moreover, we consider our estimates to be highly conservative given the fact that conservation translocations describing unsuccessful or uncertain outcomes are less likely to be published than those with successful outcomes (Fischer and Lindenmayer, 2000; Miller *et al*., 2014; Reading *et al*., 1997).

## IV. DISCUSSION

Our meta-analysis gauged the efficacy of conservation translocation programs for benefitting population and species persistence based on direct, *in situ* comparisons of animal performance between translocated organisms and wild-resident conspecifics. We synthesized estimates of relative performance based on five fitness-surrogate categories across 101 unique species spanning nine taxonomic classes. Our findings indicate that translocated organisms tend to under-perform relative to their wild counterparts, and that this pattern is largely independent of taxonomic class or the type of fitness surrogate measured. This supports previous claims for the existence of general issues across ACTs and suggests that translocated organisms are often at a major disadvantage following release (see the Introduction; e.g., Snyder *et al*., 1996; Fischer & Lindenmayer, 2000; Pérez *et al*., 2012; Germano *et al*., 2015; Sullivan, Nowak & Kwiatkowski, 2015). Under-performance of translocated cohorts was also consistent regardless of translocation strategy or the amount of captive ancestry, with some caveats. Finally, post-release performance was markedly improved in translocated cohorts exposed to pre-release enrichment relative to unenriched cohorts. Here we explore the implications of our findings and make recommendations for future translocation program assessment and monitoring procedures.

### (1) Overall model

The unexpected lack of variation in effect sizes across taxonomic classes hints at a general influence of ACTs on animals, which is in direct odds with the large proportion of study heterogeneity explained by taxonomic rank (Table S5 in Appendix S2). The high taxonomic signal (64.29%) indicates that taxonomic position explains a majority of variation in effect sizes, which would lead us to expect large differences in relative performance values among higher taxonomic groups. For example, Dochtermann *et al*., (2019) observed a consistent pattern in a phylogenetic meta-analysis quantifying behavioral heritability: little difference in effect size among species and a low phylogenetic signal. However, we see the proportion of heterogeneity explained by individual taxonomic tiers is consistently low, which could explain the minimal differentiation in effect sizes at the level of taxonomic class. Equal variances estimated at each taxonomic level is consistent with an explanation of constant phylogenetic inertia across the phylogeny, in which equal branch lengths of the phylogeny coincide with equal taxonomic branch lengths. Alternatively, if phylogenetic and taxonomic branching patterns do not coincide, then the observed results would imply variation in taxonomic inertia across evolutionary time (Hadfield & Nakagawa, 2010). To further explore these alternative explanations, and investigate patterns of correlated evolution with respect to performance of translocated animals, comparative studies focusing on biological patterns at the broadest taxonomic scales could adopt a mixed phylogenetic and taxonomic approach until an appropriate phylogeny with empirically-derived branch lengths can be resolved for all species in the data (Lynch, 1991; Hadfield & Nakagawa, 2010; Johnson *et al*., 2018).

Our meta-analysis highlights a clear bias in taxonomic representation among ACTs, with most studies involving charismatic species or those with economically important fisheries. Most notably, there is a significant lack of ACTs involving amphibians that meet our inclusion criteria. Amphibian populations are experiencing global declines and will likely require human interventions in the form of ACTs to avoid extirpation (Wake & Vredenburg, 2008; Germano & Bishop, 2009; Scheele *et al*., 2021). Recent quantitative reviews describe a positive trend in the success of amphibian translocations relative to earlier benchmarks (Dodd & Seigel, 1992; Germano & Bishop, 2009). However, these efforts fail to consider comparisons among translocated and wild-resident conspecifics as a potential index for project feasibility. Recent amphibian studies using this comparative approach have provided novel insights regarding the multi-generational influence of ACTs as well as potential pitfalls of cryopreservation techniques on ACT feasibility (Cayuela *et al*., 2019; Poo *et al*., 2022). Future assessments of translocation success should include simultaneous monitoring of a wild-control group from the donor and/or recipient populations as an additional criterion for determining project feasibility and success.

Our primary model showed high consistency in effect sizes across fitness surrogates, indicating that phenotypic traits considered as correlates of overall fitness have the potential to characterize performance differences among translocated and wild-resident cohorts. We initially predicted that fitness correlates would exhibit less consistent disparities across animal cohorts since fitness correlates are less reliable proxies of fitness than fitness components (Hunt & Hodgson, 2010). One explanation for the observed pattern is that fitness surrogates can vary in how well they reflect overall individual fitness based on the relative length and positioning of the measurement interval within the organism’s lifespan (Pekkala *et al*., 2011; Hendry *et al*., 2018). Additionally, the lack of differences in fitness surrogate effects may be due to unaccounted-for differences in life-history strategies within and among study systems. Life-history theory suggests that the uneven allocation of finite resources results in life-history trade-offs among fitness components such that maximizing one fitness component can decrease overall lifetime reproductive success (e.g., increased annual fecundity decreases annual survival; Haave-Audet, Besson, Nakagawa, & Mathot, 2022; Moyes *et al*., 2009; Roff, 2002; van Noordwijk & de Jong, 1986). Future ACT studies and meta-analyses on the topic should incorporate multiple fitness components at different life stages to assess relative quality of released organisms, such that the covariance among fitness components and thus the presence of fitness trade-offs or differing life-history strategies can be tested at the within-study level (Fincke & Hadrys, 2001; Hendry *et al*., 2018).

We were not able to evaluate sex-by-fitness surrogate interactions in the full dataset because so few studies reported sex-specific effects. However, sufficient data from reptile studies suggest that estimates of relative performance based on movement were substantially lower for both male- and female-only cohorts relative to estimates from mixed-sex cohorts, indicating greater disparity in post-release performance relative to wild conspecifics when males and females were analyzed separately. Only when accounting for sex do our findings align with the established relationship between maladaptive post-release movement patterns and reduced survival in the herpetofaunal literature (Bonnet, Naulleau & Shine, 1999; Germano & Bishop, 2009). The influence of sex on post-release performance is also poorly represented in the primary literature and rarely synthesized in subsequent reviews (Teixeira *et al*., 2007). To our knowledge, no other quantitative literature review or meta-analysis examining translocation efficacy has included sex as a moderating factor, despite well-documented and systematic differences in life history and ecology among the sexes (Immonen *et al*., 2018; Tarka *et al*., 2018).

### (2) Influence of captive ancestry

We observed similar levels of under-performance among translocated cohorts relative to wild conspecifics, regardless of whether translocated organisms were ISR (0 captive generations), head-started (ESR with < 1 captive generation), or captive-bred (ESR with > 1 captive generation). This contrasts with existing quantitative reviews describing a negative relationship between captive ancestry and fitness, with ISRs performing relatively better than ESRs (Fischer & Lindenmayer, 2000; Rummel *et al*., 2016). One explanation for this discrepancy is the poor reporting in the literature of ESR captive histories and animal origins which precluded us from conducting a more powerful assessment of the effect of captive ancestry. As such, our head-starting vs. captive-rearing levels possibly misclassified cohorts with ambiguous or mixed captive histories, which could explain the lack of significant performance deficits commonly reported in long-term captive-breeding populations (Araki *et al*., 2007). It should also be noted that the ACT strategies specific to certain taxonomic groups might be more prone to reporting ambiguous captive ancestry information (Table S9). For example, reptilian, mammalian, and avian studies regularly describe captive ancestries in great detail (Jones *et al*., 2002), whereas tracking individual captive-ancestries in fisheries systems is impossible without genetic parentage assignment (Lutz *et al*., 2021). This can be especially problematic if there is captive ancestry in the purported wild-resident group, which would deflate expected fitness differences between translocated and wild-resident conspecifics (Araki *et al*., 2007). Our findings suggest that while ISRs avoid the maladaptive selective pressures associated with extended captivity, managers should still be wary of individual- and population-level stressors characteristic of ISRs that can lead to disparities in post-release performance like those observed in ESRs. This underscores the necessity of ACT programs to report the capture histories of their organisms, and to conduct multi-generational monitoring to ensure accurate measures of meaningful life-history traits.

### (3) Influence of pre-release enrichment

Disparity in post-release performance among translocated and wild individuals was significantly reduced when translocated organisms were exposed to some form of pre-release enrichment. This finding highlights the growing relevance of the behavioral ramifications of *in situ* and *ex situ* reinforcements, where patterns of allelic heterozygosity and population growth rate have historically been the primary factors used to assess translocation efficacy (Frankham, 2008; Greggor *et al*., 2016; Mathews *et al*., 2005; Morris *et al*., 2021). Our findings agree with those from other meta-analyses showing that enriched, translocated organisms generally had higher performance than non-enriched, translocated conspecifics in a variety of taxa (Tetzlaff *et al*., 2019a; Zhang *et al*., 2022). Since few studies incorporating pre-release enrichment also fulfilled our inclusion criteria by directly comparing translocated cohorts to wild-resident conspecifics, we were not able to refine our categorization of different enrichment strategies. Despite that, our study provides the broadest evidence yet—and the first synthesized evidence to incorporate wild-control groups—that pre-release enrichment strategies have the potential to reduce post-release performance disparities between translocated and wild-resident organisms.

### (4) Future directions

Despite notable positive outcomes, animal conservation translocations have experienced historically low success rates without significant improvement observed over time (Berger-Tal, Blumstein & Swaisgood, 2020). Ensuring future success of ACTs requires improved alignment between the specific conservation benefits sought in the translocation (e.g., as defined in Annex 2 of IUCN/SSC, 2013) coupled with objective evaluation of success achieving the desired benefits. Conservation biology is in the midst of a revolutionary push for more evidence-based decision-making to replace historically anecdotal- or experience-driven strategies (Pullin & Knight, 2001; Sutherland *et al*., 2004). Formal meta-analysis is an important tool for synthesizing existing literature and directing new veins of inquiry, particularly in the field of conservation, where translocation studies are numerous but statistical power is limited by low sample sizes (Moseby, Hill & Lavery, 2014). Here we discuss several aspects of ACT practices and conservation goals that are relevant to increasing insight from future meta-analyses.

The conservation benefits of many ACTs (*sensu* IUCN/SSN, 2013) target population persistence and preventing outright species extinction. Short-term strategies for achieving these goals can be generalized according to three different ways to ‘rescue’ populations (Carlson *et al*., 2014): increase population density and buffer against ecological effects and demographic stochasticity (i.e., demographic rescue), increase genetic heterozygosity and reduce the accumulation of deleterious recessive alleles (i.e., genetic rescue), or increase adaptive potential (i.e., evolutionary rescue). Ultimately, to achieve these goals, translocation activities must contribute to populations in ways that enable recipient populations to adapt to their environment, increase average individual fitness, and thereby bolster population growth rate (Lande 1988; Carlson *et al*., 2014; Hendry *et al*., 2018; Shaw, 2019). To evaluate the density-dependent ecological dynamics associated with demographic rescue, the recipient population cannot serve as the control group and so an allopatric population should be used instead. In our systematic review, we found only one study that included both sympatric and allopatric control groups (Molony *et al*., 2006). Thus, our results can only be interpreted in light of how successfully ACTs contribute to increasing mean fitness of the recipient population.

Our findings show that *in situ* reinforcements (ISRs) and *ex situ* reinforcements (ESRs) produce organisms that under-perform and should therefore be implemented only after careful consideration of the recipient population’s trajectory and the existing environmental threats at the recipient site (IUCN/SSC, 2013). For example, if a wild population is declining due to extrinsic threats, then the release of under-performing translocated conspecifics is unlikely to reverse declines (Heppell *et al*., 1996). Alternatively, if environmental threats have been removed, then the allelic diversity of a remnant wild population could benefit from the influx of translocated cohorts, despite certain performance deficits (Jensen *et al*., 2018; Cayuela *et al*., 2019). Pilot studies employing the post-release comparative techniques analyzed in this study could therefore use relative post-release performance, population demographics, and environmental forecasts to predict the efficacy of ACT programs.

The initial prevention and reversal of invasive species spread is a major aspect of global habitat restoration which shares a conceptual basis with conservation translocations (Bright & Smithson, 2001; Armstrong & Seddon, 2008) and thus stands to benefit from *in situ* comparisons with wild-control groups. The potential for non-invasive species establishment and persistence can be assessed through sympatric comparisons with non-conspecifics of overlapping ecological niche, or through allopatric comparisons with conspecifics from the original donor population. If an invasion is ongoing, direct comparisons between recent invaders and the offspring of previous-generation invaders could also promote the investigation of the genetic basis of invasive adaptations. Similar investigations could act as the prelude to assisted colonization or rewilding efforts to ensure that species introduced to areas outside of their historic range are capable of colonizing successfully without risk of disrupting existing ecosystems (Corlett, 2016; Hunter-Ayad *et al*., 2021).

The often-wide credible intervals in our findings suggest there are opportunities in which ACTs can produce viable individuals and contribute to recipient populations. We recommend that ACT managers include sympatric and allopatric wild-reference groups in their programs to properly evaluate post-release performance of translocated organisms, and assess any negative density-dependent impacts of ACTs on resident conspecifics (Mathews *et al*., 2005; Molony *et al*., 2006). Managers should also take advantage of next-generation sequencing technologies and parentage assignment software to promote long-term, multi-generational monitoring of relative reproductive success and other fitness surrogates (Araki *et al*., 2007).

## V. CONCLUSIONS

1. Translocated organisms generally exhibited poorer performance following release compared to wild-resident conspecifics. The translocated organisms showed an overall 64% decreased odds of out-performing their wild-resident counterparts (odds ratio posterior mode: 0.30; 95% CrI: 0.18 to 0.66) and this degree of disparity was consistent across taxonomic classes as well as whether an *in situ versus ex situ* reinforcement (ISR vs. ESR) strategy had been implemented.
2. The overall disparity between translocated and wild-resident cohorts was consistent across studies regardless of the amount of captive ancestry in any particular implementation.
3. Post-release performance of translocated organisms generally benefitted as a result of some level of enrichment activity incorporated into the translocation program.
4. The dataset exhibited a high overall taxonomic signal but low heterogeneity at higher individual taxonomic levels. Variation across taxonomic levels suggests a potential influence of shared evolutionary history on relative post-release performance, but more comprehensive taxonomic sampling coupled with comparative analyses in a phylogenetic framework is necessary to draw accurate conclusions.
5. Overall, the disparity in post-release performance between translocated organisms and their wild-resident counterparts was not affected by sex, taxonomic group, type of fitness surrogate measured, or ESR vs. ISR translocation strategy.
6. Only 2% of animal conservation translocation studies returned during the initial literature search of our systematic review report or design translocations that use wild-resident control groups. We suggest both sympatric and allopatric wild-reference groups will further enable post-release performance of translocated organisms to be evaluated for any negative density-dependent impacts of ACTs on resident conspecifics.
7. Conservation managers should standardize the use of wild-resident controls and multiple performance metrics that capture several different aspects of the life history in their assessments of potential conservation translocation strategies to ensure population or species persistence.

## VI. ACKNOWLEDGEMENTS

We would like to thank members of the D.A. Warner and M.E. Wolak labs, and students from the Auburn University meta-analysis course (taught by A.E.W.) for providing comments on earlier drafts of the manuscript.

## VI. AUTHOR CONTRIBUTIONS

**Conceptualization:** IPG and MEW; **Data curation:** IPG and MEW; **Formal analysis:** IPG, with input from AEW and MEW; **Investigation:** IPG; **Project administration:** MEW; **Visualization:** IPG and MEW; **Writing - original manuscript:** IPG; **Writing - review and editing:** AEW and MEW.

## VII. DATA ACCESSIBILITY

All data and R code are available from the Zenodo Digital Repository https://doi.org/10.5281/zenodo.7535746

## X. SUPPORTING INFORMATION

Additional supporting information may be found online in the Supporting Information section at the end of the article.

**Appendix S1:** PRISMA EcoEvo checklist.

**Appendix S2:** Supplemental tables summarizing full-text screening outcomes, meta-analysis model summary tables, and heterogeneity statistics.

**Appendix S3:** Supplemental figures.

## REFERENCES

Aaltonen, K., Bryant, A.A., Hostetler, J.A. & Oli, M.K. (2009) Reintroducing endangered Vancouver Island marmots: Survival and cause-specific mortality rates of captive-born versus wild-born individuals. Biological Conservation 142, 2181–2190.

Aarestrup, K., Baktoft, H., Koed, A., Del Villar-Guerra, D. & Thorstad, E. (2014) Comparison of the riverine and early marine migration behaviour and survival of wild and hatchery-reared sea trout Salmo trutta smolts. Marine Ecology Progress Series 496, 197–206.

Agnalt, A.L., Kristiansen, T.S. & Jørstad, K.E. (2007) Growth, reproductive cycle, and movement of berried European lobsters (*Homarus gammarus*) in a local stock off southwestern Norway. ICES Journal of Marine Science 64, 288–297.

Aguinis, H., Gottfredson, R.K. & Joo, H. (2013) Best-practice recommendations for defining, identifying, and handling outliers. Organizational Research Methods 16, 270–301.

Alleaume-Benharira, M., Pen, I.R. & Ronce, O. (2006) Geographical patterns of adaptation within a species’ range: Interactions between drift and gene flow. Journal of Evolutionary Biology 19, 203–215.

Anderson, J.H., Faulds, P.L., Atlas, W.I. & Quinn, T.P. (2013) Reproductive success of captively bred and naturally spawned Chinook salmon colonizing newly accessible habitat. Evolutionary Applications 6, 165–179.

Araki, H., Berejikian, B.A., Ford, M.J. & Blouin, M.S. (2008) Fitness of hatchery-reared salmonids in the wild. Evolutionary Applications 1, 342–355.

Araki, H., Cooper, B. & Blouin, M.S. (2007) Genetic effects of captive breeding cause a rapid, cumulative fitness decline in the wild. Science 318, 100–103.

Armstrong, D.P. & Seddon, P.J. (2008) Directions in reintroduction biology. Trends in Ecology and Evolution 23, 20–25.

Attum, O. & Cutshall, C.D. (2015) Movement of translocated turtles according to translocation method and habitat structure. Restoration Ecology 23, 588–594.

Bacon, L., Robert, A. & Hingrat, Y. (2019) Long lasting breeding performance differences between wild-born and released females in a reinforced North African houbara bustard (*Chlamydotis undulata undulata*) population: a matter of release strategy. Biodiversity and Conservation 28, 553–570.

Bajomi, B., Pullin, A.S., Stewart, G.B. & Takács-Sánta, A. (2010) Bias and dispersal in the animal reintroduction literature. Oryx 44, 358–365.

Barnosky, A.D., Matzke, N., Tomiya, S., Wogan, G.O.U., Swartz, B., Quental, T.B., Marshall, C., Mcguire, J.L., Lindsey, E.L., Maguire, K.C., Mersey, B. & Ferrer, E.A. (2011) Has the Earth’s sixth mass extinction already arrived? Nature 471, 51–57.

Bauder, J.M., Castellano, C., Jensen, J.B., Stevenson, D.J. & Jenkins, C.L. (2014) Comparison of movements, body weight, and habitat selection between translocated and resident gopher tortoises. The Journal of Wildlife Management 78, 1444–1455.

Bennett, A.M., Steiner, J., Carstairs, S., Gielens, A. & De, C.M.D. (2017) A question of scale: replication and the effective evaluation of conservation interventions. Facets 2, 892– 909.

Berger-Tal, O., Blumstein, D.T. & Swaisgood, R.R. (2020) Conservation translocations: a review of common difficulties and promising directions. Animal Conservation 23, 121–131.

Beringer, J., Hansen, L.P., Demand, J.A., Sartwell, J., Wallendorf, M. & Mange, R. (2002) Efficacy of translocation to control urban deer in Missouri: costs, efficiency, and outcome. Wildlife Society Bulletin 30, 767–774.

Blanchard, B.M. & Knight, R.R. (1995) Biological consequences of relocating grizzly bears in the Yellowstone ecosystem. The Journal of Wildlife Management 59, 560–565.

Blythe, R.M., Smyser, T.J., Johnson, S.A. & Swihart, R.K. (2015) Post-release survival of captive-reared Allegheny woodrats. Animal Conservation 18, 186–195.

Bolam, F.C., Mair, L., Angelico, M., Brooks, T.M., Burgman, M., Hermes, C., Hoffmann, M., Martin, R.W., Mcgowan, P.J.K., Rodrigues, A.S.L., Rondinini, C., Westrip, J.R.S., Wheatley, H., Bedolla-Guzmán, Y., Calzada, J., et al. (2021) How many bird and mammal extinctions has recent conservation action prevented? Conservation Letters 14, e12762.

Bolland, J.D., Cowx, I.G. & Lucas, M.C. (2009) Dispersal and survival of stocked cyprinids in a small English river: comparison with wild fishes using a multi-method approach. Journal of Fish Biology 74, 2313–2328.

Bonduriansky, R. & Chenoweth, S.F. (2009) Intralocus sexual conflict. Trends in Ecology and Evolution 24, 280–288.

Bonnet, X., Naulleau, G. & Shine, R. (1999) The dangers of leaving home: dispersal and mortality in snakes. Biological Conservation 89, 39–50.

Borgstrom, R., Skaala, Ø. & Aastveit, A.H. (2002) High mortality in introduced brown trout depressed potential gene flow to a wild population. Journal of Fish Biology 61, 1085– 1097.

Bose, A.P.H., Zayonc, D., Avrantinis, N., Ficzycz, N., Fischer-Rush, J., Francis, F.T., Gray, S., Manning, F., Robb, H., Schmidt, C., Spice, C., Umedaly, A., Warden, J. & Côté, I.M. (2019) Effects of handling and short-term captivity: a multi-behaviour approach using red sea urchins, *Mesocentrotus franciscanus*. PeerJ 7, e6556.

Bosson, C.O., Palme, R. & Boonstra, R. (2013) Assessing the impact of live-capture, confinement, and translocation on stress and fate in eastern gray squirrels. Journal of Mammalogy 94, 1401–1411.

Bourass, K. & Hingrat, Y. (2015) Diet of released captive-bred North-African houbara bustards. European Journal of Wildlife Research 61, 563–574.

Bouzat, J.L., Johnson, J.A., Toepfer, J.E., Simpson, S.A., Esker, T.L. & Westemeier, R.L. (2009) Beyond the beneficial effects of translocations as an effective tool for the genetic restoration of isolated populations. Conservation Genetics 10, 191–201.

Bradley, E.H., Pletscher, D.H., Bangs, E.E., Kunkel, K.E., Smith, D.W., Mack, C.M., Meier, T.J., Fontaine, J.A., Niemeyer, C.C. & Jimenez, M.D. (2005) Evaluating translocation as a nonlethal method to reduce livestock conflicts in the northwestern United States. Conservation Biology 19, 1498–1508.

Brand, L.A., Farnsworth, M.L., Meyers, J., Dickson, B.G., Grouios, C., Scheib, A.F. & Scherer, R.D. (2016) Mitigation-driven translocation effects on temperature, condition, growth, and mortality of Mojave desert tortoise (*Gopherus agassizii*) in the face of solar energy development. Biological Conservation 200, 104–111.

Brannelly, L.A., Hunter, D.A., Skerratt, L.F., Scheele, B.C., Lenger, D., Mcfadden, M.S., Harlow, P.S. & Berger, L. (2016) Chytrid infection and post-release fitness in the reintroduction of an endangered alpine tree frog. Animal Conservation 19, 153–162.

Brichieri-Colombi, T.A., Lloyd, N.A., Mcpherson, J.M. & Moehrenschlager, A. (2019) Limited contributions of released animals from zoos to North American conservation translocations. Conservation Biology 33, 33–39.

Bright, P.W. & Smithson, T.J. (2001) Biological invasions provide a framework for reintroductions: Selecting areas in England for pine marten releases. Biodiversity and Conservation 10, 1247–1265.

Brittas, R., Marcström, V., Kenward, R.E. & Karlbom, M. (1992) Survival and breeding success of reared and wild ring-necked pheasants in Sweden. The Journal of Wildlife Management 56, 368–376.

Brown, C., Davidson, T. & Laland, K. (2003) Environmental enrichment and prior experience of live prey improve foraging behaviour in hatchery-reared Atlantic salmon. Journal of Fish Biology 63, 187–196.

Brown, J.R., Bishop, C.A. & Brooks, R.J. (2009) Effectiveness of short-distance translocation and its effects on western rattlesnakes. The Journal of Wildlife Management 73, 419–425.

Brown, J.L., Collopy, M.W., Gott, E.J., Juergens, P.W., Montoya, A.B. & Hunt, W.G. (2006) Wild-reared aplomado falcons survive and recruit at higher rates than hacked falcons in a common environment. Biological Conservation 131, 453–458.

Burnside, R.J., Collar, N.J. & Dolman, P.M. (2017) Comparative migration strategies of wild and captive-bred Asian houbara *Chlamydotis macqueenii*. Ibis 159, 374–389.

Burnside, R.J., Collar, N.J. & Dolman, P.M. (2018) Dataset on the numbers and proportion of mortality attributable to hunting, trapping, and powerlines in wild and captive-bred migratory Asian houbara *Chlamydotis macqueenii*. Data in Brief 21, 1848–1852.

Butler, H., Malone, B. & Clemann, N. (2005) The effects of translocation on the spatial ecology of tiger snakes (*Notechis scutatus*) in a suburban landscape. Wildlife Research 32, 165.

Carlson, S.M., Cunningham, C.J. & Westley, P.A.H. (2014) Evolutionary rescue in a changing world. Trends in Ecology & Evolution 29, 521–530.

Cayuela, H., Gillet, L., Laudelout, A., Besnard, A., Bonnaire, E., Levionnois, P., Muths, E., Dufrêne, M. & Kinet, T. (2019) Survival cost to relocation does not reduce population self-sustainability in an amphibian. Ecological Applications 29, 1–15.

Ceballos, G., Ehrlich, P.R., Barnosky, A.D., García, A., Pringle, R.M. & Palmer, T.M. (2015) Accelerated modern human–induced species losses: entering the sixth mass extinction. Science Advances 1, e1400253.

Ceballos, G., Ehrlich, P.R. & Dirzo, R. (2017) Biological annihilation via the ongoing sixth mass extinction signaled by vertebrate population losses and declines. Proceedings of the National Academy of Sciences of the United States of America 114, E6089–E6096.

Ceballos, G., Ehrlich, P.R. & Raven, P.H. (2020) Vertebrates on the brink as indicators of biological annihilation and the sixth mass extinction. Proceedings of the National Academy of Sciences of the United States of America 117, 13596–13602.

Champagnon, J., Guillemain, M., Elmberg, J., Massez, G., Cavallo, F. & Gauthier-Clerc, M. (2012) Low survival after release into the wild: assessing “the burden of captivity” on mallard physiology and behaviour. European Journal of Wildlife Research 58, 255–267.

Chinn, S. (2000) A simple method for converting an odds ratio to effect size for use in metalJanalysis. Statistics in Medicine 19, 3127–3131.

Chittaro, P., Johnson, L., Teel, D., Moran, P., Sol, S., Macneale, K. & Zabel, R. (2018) Variability in the performance of juvenile Chinook salmon is explained primarily by when and where they resided in estuarine habitats. Ecology of Freshwater Fish 27, 857–873.

Chittenden, C.M., Sura, S., Butterworth, K.G., Cubitt, K.F., Plantalech Manel-la, N., Balfry, S., Økland, F. & Mckinley, R.S. (2008) Riverine, estuarine and marine migratory behaviour and physiology of wild and hatchery-reared coho salmon *Oncorhynchus kisutch* (Walbaum) smolts descending the Campbell River, Bc, Canada. Journal of Fish Biology 72, 614–628.

Clifford, D.L., Woodroffe, R., Garcelon, D.K., Timm, S.F. & Mazet, J. A. K. (2007) Using pregnancy rates and perinatal mortality to evaluate the success of recovery strategies for endangered island foxes. Animal Conservation 10, 442–451.

Clutton-Brock, T.H. & Harvey, P.H. (1977) Primate ecology and social organization. Journal of Zoology 183, 1–39.

Cohen, J. (1992) Quantitative methods in psychology: a power primer. Psychological Bulletin 112, 1155–1159.

Collis, K., Roby, D.D., Craig, D.P., Ryan, B.A. & Ledgerwood, R.D. (2001) Colonial waterbird predation on juvenile salmonids tagged with passive integrated transponders in the Columbia River estuary: vulnerability of different salmonid species, stocks, and rearing types. Transactions of the American Fisheries Society 130, 385–396.

Corlett, R.T. (2016) Restoration, reintroduction, and rewilding in a changing world. Trends in Ecology and Evolution 31, 453–462.

Coulson, T., Benton, T.G., Lundberg, P., Dall, S.R.X., Kendall, B.E. & Gaillard, J.M. (2006) Estimating individual contributions to population growth: evolutionary fitness in ecological time. Proceedings of the Royal Society B: Biological Sciences 273, 547–555.

Crates, R., Stojanovic, D. & Heinsohn, R. (2022) The phenotypic costs of captivity. Biological Reviews, 434–449.

Cromwell, J.A., Warren, R.J. & Henderson, D.W. (1999) Live-capture and small-scale relocation of urban deer on Hilton Head Island, South Carolina. Wildlife Society Bulletin (1973-2006) 27, 1025–1031.

Dannewitz, J., Petersson, E., Dahl, J., Prestegaard, T., Löf, A.-C. & Järvi, T. (2004) Reproductive success of hatchery-produced and wild-born brown trout in an experimental stream. Journal of Applied Ecology 41, 355–364.

Davis, J.L.D., Eckert-Mills, M.G., Young-Williams, A.C., Hines, A.H. & Zohar, Y. (2005) Morphological conditioning of a hatchery-raised invertebrate, *Callinectes sapidus*, to improve field survivorship after release. Aquaculture 243, 147–158.

Davis, J.L.D., Young-Williams, A.C., Aguilar, R., Carswell, B.L., Goodison, M.R., Hines, A.H., Kramer, M.A., Zohar, Y. & Zmora, O. (2004) Differences between hatchery-raised and wild blue crabs: implications for stock enhancement potential. Transactions of the American Fisheries Society 133, 1–14.

Davis, M.J., Woo, I., Ellings, C.S., Hodgson, S., Beauchamp, D.A., Nakai, G. & De La Cruz, S.E.W. (2018) Integrated diet analyses reveal contrasting trophic niches for wild and hatchery juvenile chinook salmon in a large river delta. Transactions of the American Fisheries Society 147, 818–841.

Davis, M.J., Woo, I., Ellings, C.S., Hodgson, S., Beauchamp, D.A., Nakai, G. & De La Cruz, S.E.W. (2019) Freshwater tidal forests and estuarine wetlands may confer early life growth advantages for delta-reared chinook salmon. Transactions of the American Fisheries Society 148, 289–307.

Day, T. & Bonduriansky, R. (2011) A unified approach to the evolutionary consequences of genetic and nongenetic inheritance. American Naturalist 178.

Degregorio, B.A., Sperry, J.H., Tuberville, T.D. & Weatherhead, P.J. (2017) Translocating ratsnakes: Does enrichment offset negative effects of time in captivity? Wildlife Research 44, 438–448.

Degregorio, B.A., Weatherhead, P.J., Tuberville, T.D. & Sperry, J.H. (2013) Time in captivity affects foraging behavior of ratsnakes: implications for translocation. Herpetological Conservation and Biology 8, 581–590.

Del Real, S.C., Workman, M. & Merz, J. (2012) Migration characteristics of hatchery and natural-origin *Oncorhynchus mykiss* from the lower Mokelumne River, California. Environmental Biology of Fishes 94, 363–375.

Delgado, G.A., Bartels, C.T., Glazer, R.A., Brown-Peterson, N.J. & Mccarthy, K.J. (2004) Translocation as a strategy to rehabilitate the queen conch (*Strombus gigas*) Population in the Florida Keys. Fishery Bulletin 102, 278–288.

Devan-Song, A., Martelli, P., Dudgeon, D., Crow, P., Ades, G. & Karraker, N.E. (2016) Is long-distance translocation an effective mitigation tool for white-lipped pit vipers (*Trimeresurus albolabris*) in South China? Biological Conservation 204, 212–220.

Dickens, M.J., Delehanty, D.J. & Michael Romero, L. (2010) Stress: An inevitable component of animal translocation. Biological Conservation 143, 1329–1341.

Dochtermann, N.A., Schwab, T., Anderson Berdal, M., Dalos, J. & Royauté, R. (2019) The heritability of behavior: a meta-analysis. Journal of Heredity 110, 403–410.

Dodd, C.K. & Seigel, R.A. (1992) Relocation, repatriation, and translocation of amphibians and reptiles: are they conservation strategies that work? Biological Conservation 62, 230.

Drake, K.K., Nussear, K.E., Esque, T.C., Barber, A.M., Vittum, K.M., Medica, P.A., Tracy, C.R. & Hunter Jr, K.W. (2012) Does translocation influence physiological stress in the desert tortoise? Animal Conservation 15, 560–570.

Ducatez, S. & Shine, R. (2019) Life-history traits and the fate of translocated populations. Conservation Biology 33, 853–860.

Duval, S. & Tweedie, R. (2000) Trim and fill: a simple funnel-plot-based method. Biometrics 56, 455–463.

Egger, M., Smith, G.D., Schneider, M. & Minder, C. (1997) Bias in meta-analysis detected by a simple, graphical test. British Medical Journal 315, 629–634.

Elsey, R.M., Joanen, T., Mcnease, L. & Kinler, N. (1992) Growth rates and body condition factors of *Alligator mississippiensis* in coastal Louisiana wetlands: A comparison of wild and farm-released juveniles. Comparative biochemistry and physiology. Part A, Molecular & integrative physiology 103a, 667–672.

Elsey, R.M., Mcnease, L. & Joanen, T. (2000) Louisiana’s alligator ranching programme: a review and analysis of releases of captive-raised juveniles. In Crocodilian Biology and Evolution pp. 426–441. Surrey Beatty & Sons, Chipping Norton.

Escobar, R.A., Besier, E. & Hayes, W.K. (2010) Evaluating headstarting as a management tool: post-release success of green iguanas (*Iguana iguana*) in Costa Rica. International Journal of Biodiversity and Conservation 2, 204–214.

Esque, T., Nussear, K., Drake, K., Walde, A., Berry, K., Averill-Murray, R., Woodman, A., Boarman, W., Medica, P., Mack, J. & Heaton, J. (2010) Effects of subsidized predators, resource variability, and human population density on desert tortoise populations in the Mojave Desert, USA. Endangered Species Research 12, 167–177.

Evans, A.F., Hostetter, N.J., Collis, K., Roby, D.D. & Loge, F.J. (2014) Relationship between Juvenile Fish Condition and Survival to Adulthood in Steelhead. Transactions of the American Fisheries Society 143, 899–909.

Evans, M.L., Johnson, M.A., Jacobson, D., Wang, J., Hogansen, M. & O’malley, K.G. (2016) Evaluating a multi-generational reintroduction program for threatened salmon using genetic parentage analysis. Canadian Journal of Fisheries and Aquatic Sciences 73, 844– 852.

Evans, M.L., Wilke, N.F., O’reilly, P.T. & Fleming, I.A. (2014) Transgenerational effects of parental rearing environment influence the survivorship of captive-born offspring in the wild. Conservation Letters 7, 371–379.

Fairbairn, D.J. (2013) Odd couples: Extraordinary differences between the sexes in the animal kingdom. Princeton University Press.

Fairbairn, D.J. & Reeve, J.P. (2001) Natural selection: Measuring selection in natural populations. In Evolutionary Ecology: Concepts and case studies (eds C. Fox, D.A. Roff & D.J. Fairbairn), Oxford University Press, NY.

Farnsworth, M.L., Dickson, B.G., Zachmann, L.J., Hegeman, E.E., Cangelosi, A.R., Jackson, T.G. & Scheib, A.F. (2015) Short-term space-use patterns of translocated Mojave Desert tortoise in Southern California. PLOS ONE 10, e0134250.

Farnsworth, S.D. & Seigel, G., Richard A. (2013) Responses, movements, and survival of relocated box turtles during the construction of the Inter-County Connector Highway in Maryland. Transportation Research Record Journal of the Transportation Research Board 2362, 1–8.

Farquharson, K.A., Hogg, C.J. & Grueber, C.E. (2018) A meta-analysis of birth-origin effects on reproduction in diverse captive environments. Nature Communications 9, 1–10.

Farquharson, K.A., Hogg, C.J. & Grueber, C.E. (2021) Offspring survival changes over generations of captive breeding. Nature Communications 12, 1–9.

Fernando, P., Leimgruber, P., Prasad, T. & Pastorini, J. (2012) Problem-elephant translocation: translocating the problem and the elephant? PLoS ONE 7, e50917.

Ferretti, M., Paci, G., Porrini, S., Galardi, L. & Bagliacca, M. (2010) Habitat use and home range traits of resident and relocated hares (*Lepus europaeus*, Pallas). Italian Journal of Animal Science 9, e54.

Ficetola, G.F. & DE Bernardi, F. (2005) Supplementation or *in situ* conservation? Evidence of local adaptation in the Italian agile frog *Rana latastei* and consequences for the management of populations. Animal Conservation 8, 33–40.

Finstad, B. & Heggberget, T.G. (1993) Migration, growth and survival of wild and hatchery-reared anadromous Arctic charr (*Salvelinus alpinus*) in Finnmark, Northern Norway. Journal of Fish Biology 43, 303–312.

Fincke, O.M. & Hadrys, H. (2001) Unpredictable offspring survivorship in the damselfly, *megaloprepus coerulatus*, shapes parental behavior, constrains sexual selection, and challenges traditional fitness estimates. Evolution 55, 762–772.

Fischer, J. & Lindenmayer, D.B. (2000) An assessment of the published results of animal relocations. Biological Conservation 96, 1–11.

Fjellheim, A., Raddum, G.G. & Barlaup, B.T. (1995) Dispersal, growth and mortality of brown trout (*Salmo trutta L*.) stocked in a regulated west Norwegian river. Regulated Rivers: Research & Management 10, 137–145.

Flávio, H., Aarestrup, K., Jepsen, N. & Koed, A. (2019) Naturalised Atlantic salmon smolts are more likely to reach the sea than wild smolts in a lowland fjord. River Research and Applications 35, 216–223.

Foley, A.M., Pierce, B., Hewitt, D.G., Deyoung, R.W., Campbell, T.A., Hellickson, M.W., Ranch, K., Box, P.O., Feild, J., Ranch, K., Box, P.O., Mitchell, S., Parks, T., Lockwood, M.A., Parks, T., et al. (2008) Survival and movements of translocated white-tailed deer in South Texas.

Ford, M.J. (2002) Selection in captivity during supportive breeding may reduce fitness in the wild. Conservation Biology 16, 815–825.

Ford, M.J., Murdoch, A. & Howard, S. (2012) Early male maturity explains a negative correlation in reproductive success between hatchery-spawned salmon and their naturally spawning progeny. Conservation Letters 5, 450–458.

Ford, M.J., Murdoch, A.R., Hughes, M.S., Seamons, T.R. & Lahood, E.S. (2016) Broodstock history strongly influences natural spawning success in hatchery steelhead (*Oncorhynchus mykiss*). PLoS One, 1–20.

Frankham, R. (2008) Genetic adaptation to captivity in species conservation programs. Molecular Ecology 17, 325–333.

Furlan, E.M., Gruber, B., Attard, C.R.M., Wager, R.N.E., Kerezsy, A., Faulks, L.K., Beheregaray, L.B. & Unmack, P.J. (2020) Assessing the benefits and risks of translocations in depauperate species: A theoretical framework with an empirical validation. Journal of Applied Ecology 57, 831–841.

Gelman, A. (2006) Prior distributions for variance parameters in hierarchical models (comment on article by Browne and Draper). Bayesian Analysis 1, 515–534.

Germano, J.M. & Bishop, P.J. (2009) Suitability of amphibians and reptiles for translocation. Conservation Biology 23, 7–15.

Germano, J.M., Field, K.J., Griffiths, R.A., Clulow, S., Foster, J., Harding, G. & Swaisgood, R.R. (2015) Mitigation-driven translocations: are we moving wildlife in the right direction? Frontiers in Ecology and the Environment 13, 100–105.

Gilmartin, W.G., Sloan, A.C., Harting, A.L., Johanos, T.C., Baker, J.D., Breese, M. & Ragen, T.J. (2011) Rehabilitation and relocation of young Hawaiian monk seals (*Monachus schauinslandi*). Aquatic Mammals 37, 332–341.

Glass, G. V. (1976) Primary, secondary, and meta-analysis of research. Educational Researcher 5, 3–8.

Goetz, F.A., Jeanes, E., Moore, M.E. & Quinn, T.P. (2015) Comparative migratory behavior and survival of wild and hatchery steelhead (*Oncorhynchus mykiss*) smolts in riverine, estuarine, and marine habitats of Puget Sound, Washington. Environmental Biology of Fishes 98, 357–375.

Goldenberg, S.Z., Owen, M.A., Brown, J.L., Wittemyer, G., Oo, Z.M. & Leimgruber, P. (2019) Increasing conservation translocation success by building social functionality in released populations. Global Ecology and Conservation 18, e00604.

Goossen, J.P., Vander Lee, R., Kruse, C., Gratto-Trevor, C.L. & Westworth, S.M. (2011) Resightings of captive-reared and wild piping plovers from Saskatchewan, Canada. Wader Study Group Bulletin 118, 118–122.

Greggor, A.L., Berger-Tal, O., Blumstein, D.T., Angeloni, L., Bessa-Gomes, C., Blackwell, B.F., ST Clair, C.C., Crooks, K., de Silva, S., Fernández-Juricic, E., Goldenberg, S.Z., Mesnick, S.L., Owen, M., Price, C.J., Saltz, D., et al. (2016) Research priorities from animal behaviour for maximising conservation progress. Trends in Ecology and Evolution 31, 953–964.

Griffith, B., Scott, M.J., Carpenter, J.W. & Reed, C. (1989) Translocation as a species conservation tool: status and strategy. Science 245, 477–480.

Gurevitch, J., Koricheva, J., Nakagawa, S. & Stewart, G. (2018) Meta-analysis and the science of research synthesis. Nature 555, 175–182.

Haave-Audet, E., Besson, A.A., Nakagawa, S. & Mathot, K.J. (2022) Differences in resource acquisition, not allocation, mediate the relationship between behaviour and fitness: a systematic review and meta-analysis. Biological Reviews 97, 708–731.

Haddock, C.K., Rindskopf, D. & Shadish, W.R. (1998) Using odds ratios as effect sizes for meta-analysis of dichotomous data: a primer on methods and issues. Psychological Methods 3, 339–353.

Hadfield, J.D. (2010) MCMCglmm: MCMC methods for multi-response GLMMs in R. Journal of Statistical Software 33, 1–22.

Hadfield, J.D. & Nakagawa, S. (2010) General quantitative genetic methods for comparative biology: phylogenies, taxonomies and multi-trait models for continuous and categorical characters. Journal of Evolutionary Biology 23, 494–508.

Hadjisterkotis, E. (1999) The survival of captive bred chukar *Alectoris chukar cypriotes*, released for restocking in Cyprus. Zeitschrift Fur Jagdwissenshaft 45, 238–249.

Hagelin, A., Calles, O., Greenberg, L., Piccolo, J. & Bergman, E. (2016) Spawning migration of wild and supplementary stocked landlocked Atlantic salmon (*Salmo Salar*). River Research and Applications 32, 383–389.

Hameau, O. & Millon, A. (2019) Assessing the effectiveness of bird rehabilitation: temporarily captive-reared little owls (*Athene noctua*) experience a similar recruitment rate as wild birds. Journal of Ornithology 160, 581–585.

Hansen, S.C. & Gosselin, L.A. (2016) Are hatchery-reared abalone naïve of predators? Comparing the behaviours of wild and hatchery-reared northern abalone, *Haliotis kamtschatkana* (Jonas, 1845). Aquaculture Research 47, 1727–1736.

Haring, M.W., Johnston, T.A., Wiegand, M.D., Fisk, A.T. & Pitcher, T.E. (2016) Differences in egg quantity and quality among hatchery- and wild-origin Chinook salmon (*Oncorhynchus tshawytscha*). Canadian Journal of Fisheries and Aquatic Sciences 73, 737– 746.

Heath, S.R., Kershner, E.L., Cooper, D.M., Lynn, S., Turner, J.M., Warnock, N., Farabaugh, S., Brock, K. & Garcelon, D.K. (2008) Rodent control and food supplementation increase productivity of endangered San Clemente loggerhead shrikes (*Lanius ludovicianus mearnsi*). Biological Conservation 141, 2506–2515.

Heath, D.D., Heath, J.W., Bryden, C.A., Johnson, R.M. & Fox, C.W. (2003) Rapid evolution of egg size in captive Salmon. Science 299, 1738–1741.

Hellstedt, P. & Kallio, E.R. (2005) Survival and behaviour of captive-born weasels (*Mustela nivalis nivalis*) released in nature. Journal of Zoology 266, 37–44.

Hendry, A.P., Schoen, D.J., Wolak, M.E. & Reid, J.M. (2018) The contemporary evolution of fitness. Annual Review of Ecology, Evolution, and Systematics 49, 457–476.

Henriquez, M.C., Macey, S.K., Baker, E.E., Kelly, L.B., Betts, R.L., Rubbo, M.J. & Clark, J.A. (2017) Translocated and resident eastern box turtles (*Terrapene c. carolina*) in New York: movement patterns and habitat use. Northeastern Naturalist 24, 249–266.

Heppell, S.S., Crowder, L.B. & Crouse, D.T. (1996) Models to evaluate headstarting as a management tool for long-lived turtles. Ecological Applications 6, 556–565.

Hess, M.A., Rabe, C.D., Vogel, J.L., Stephenson, J.J., Nelson, D.D. & Narum, S.R. (2012) Supportive breeding boosts natural population abundance with minimal negative impacts on fitness of a wild population of Chinook salmon. Molecular Ecology 21, 5236–5250.

Hester, J.M., Price, S.J. & Dorcas, M.E. (2008) Effects of relocation on movements and home ranges of eastern box turtles. The Journal of Wildlife Management 72, 772–777.

Higgins, J.P.T. & Thompson, S.G. (2002) Quantifying heterogeneity in a meta-analysis. Statistics in Medicine 21, 1539–1558.

Hill, D. & Robertson, P. (1988) Breeding success of wild and hand-reared ring-necked pheasants. The Journal of Wildlife Management 52, 446–450.

Himes, J.G., Hardy, L.M., Rudolph, D.C. & Burgdorp, S.J. (2006) Movement patterns and habitat selection by native and repatriated Louisiana pine snakes (*Pituophis ruthveni*): implications for conservation. Herpetological Natural History 9, 103–116.

Hoffmann, M., Hilton-Taylor, C., Angulo, A., Böhm, M., Brooks, T.M., Butchart, S.H.M., Carpenter, K.E., Chanson, J., Collen, B., Cox, N.A., Darwall, W.R.T., Dulvy, N.K., Harrison, L.R., Katariya, V., Pollock, C.M., ET AL. (2010) The impact of conservation on the status of the world’s vertebrates. Science 330, 1503–1509.

Holding, M.L., Frazier, J.A., Taylor, E.N. & Strand, C.R. (2012) Experimentally altered navigational demands induce changes in the cortical forebrain of free-ranging Northern Pacific rattlesnakes *(Crotalus o. oreganus)*. *Brain*, Behavior and Evolution 79, 144–154.

Horreo, J.L., Valiente, A.G., Ardura, A., Blanco, A., Garcia-Vazquez, C. & Garcia-Gonzalez, E. (2018) Nature versus nurture? Consequences of short captivity in early stages. Ecology and Evolution 8, 521–529.

Hufbauer, R.A., Szucs, M., Kasyon, E., Youngberg, C., Koontz, M.J., Richards, C., Tuff, T. & Melbourne, B.A. (2015) Three types of rescue can avert extinction in a changing environment. Proceedings of the National Academy of Sciences 112, 10557–10562.

Hunt, J. & Hodgson, D. (2010) What is fitness, and how do we measure it? In Evolutionary Behavioral Ecology (eds D. Westneat & C. Fox), p. New York: Oxford University Press.

Hunter-Ayad, J., Jarvie, S., Greaves, G., Digby, A., Ohlemüller, R., Recio, M.R. & Seddon, P.J. (2021) Novel conditions in conservation translocations: a conservative-extrapolative strategic framework. Frontiers in Conservation Science 2, 1–14.

Huntingford, F.A. (2004) Implications of domestication and rearing conditions for the behaviour of cultivated fishes. Journal of Fish Biology 65, 122–142.

Immonen, E., Hämäläinen, A., Schuett, W. & Tarka, M. (2018) Evolution of sex-specific pace-of-life syndromes: genetic architecture and physiological mechanisms. Behavioral Ecology and Sociobiology 72, 60.

Islam, M.Z., Singh, A., Basheer, M.P., Judas, J. & Boug, A. (2013) Differences in space use and habitat selection between captive-bred and wild-born houbara bustards in Saudi Arabia: results from a long-term reintroduction program. Journal of Zoology 289, 251–261.

ITIS (2021) Retrieved 27 Sep 2021 from the Integrated Taxonomic Information System (ITIS). Integrated Taxonomic Information System (ITIS). Www.itis.gov [accessed 27 September 2021].

IUCN/SCC (2013) Guidelines for reintroductions and other cnservation translocations. Version 1.0. Gland, Switzerland: IUCN Species Survival Commission.

Jachowski, D.S., Gitzen, R.A., Grenier, M.B., Holmes, B. & Millspaugh, J.J. (2011) The importance of thinking big: large-scale prey conservation drives black-footed ferret reintroduction success. Biological Conservation 144, 1560–1566.

Jennions, M.D. & Møller, A.P. (2002) Publication bias in ecology and evolution: An empirical assessment using the ‘trim and fill’ method. Biological Reviews of the Cambridge Philosophical Society 77, 211–222.

Jensen, E.L., Edwards, D.L., Garrick, R.C., Miller, J.M., Gibbs, J.P., Cayot, L.J., Tapia, W., Caccone, A. & Russello, M.A. (2018) Population genomics through time provides insights into the consequences of decline and rapid demographic recovery through head-starting in a Galapagos giant tortoise. Evolutionary Applications, 1–11.

Johnson, M.A., Francis, C.D., Miller, E.T., Downs, C.J. & Vitousek, M.N. (2018) Detecting bias in large-scale comparative analyses: Methods for expanding the scope of hypothesis-testing with hormonebase. Integrative and Comparative Biology 58, 720–728.

Johnson, O., Neely, K. & Waples, R. (2004) Lopsided fish in the Snake River Basin – fluctuating asymmetry as a way of assessing impact of hatchery supplementation in Chinook salmon, *Oncorhynchus tshawytscha*. Environmental Biology of Fishes 69, 379–393.

Johnson, S.L., Power, J.H., Wilson, D.R. & Ray, J. (2010) A comparison of the survival and migratory behavior of hatchery-reared and naturally-reared steelhead smolts in the Alsea River and Estuary, Oregon, using acoustic telemetry. North American Journal of Fisheries Management 30, 55–71.

Jokikokko, E., Kallio-Nyberg, I., Saloniemi, I. & Jutila, E. (2006) The survival of semi-wild, wild and hatchery-reared Atlantic salmon smolts of the Simojoki River in the Baltic Sea. Journal of Fish Biology 68, 430–442.

Jones, J.M. & Witham, J.H. (1990) Post-translocation survival and movements of metropolitan white-tailed deer. Wildlife Society Bulletin 18, 434–441.

Jones, K.L., Glenn, T.C., Lacy, R.C., Pierce, J.R., Unruh, N., Mirande, C.M. & Chavez-Ramirez, F. (2002) Refining the whooping crane studbook by incorporating microsatellite DNA and leg-banding analyses. Conservation Biology 16, 789–799.

Jonsson, B., Jonsson, N. & Hansen, L.P. (1990) Does juvenile experience affect migration and spawning of adult Atlantic salmon? Behavioral Ecology and Sociobiology 26, 225–230.

Jonsson, N., Jonsson, B. & Fleming, I.A. (1996) Does early growth cause a phenotypically plastic response in egg production of Atlantic salmon? Functional Ecology 10, 89–96.

Jule, K.R., Leaver, L.A. & Lea, S.E.G. (2008) The effects of captive experience on reintroduction survival in carnivores: a review and analysis. Biological Conservation 141, 355–363.

Jutila, E., Jokikokko, E., Kallio-Nyberg, I., Saloniemi, I. & Pasanen, P. (2003) Differences in sea migration between wild and reared Atlantic salmon (*Salmo salar L*.) in the Baltic Sea. Fisheries Research 60, 333–343.

Kålås, J.A., Heggberget, T.G., Bjørn, P.A. & Reitan, O. (1993) Feeding behaviour and diet of goosanders *(Mergus merganser)* in relation to salmonid seaward migration. Aquatic Living Resources 6, 31–38.

Kallio-Nyberg, I., Romakkaniemi, A., Jokikokko, E., Saloniemi, I. & Jutila, E. (2015) Differences between wild and reared *Salmo salar* stocks of two northern Baltic Sea rivers. Fisheries Research 165, 85–95.

Kellison, G., Eggleston, D., Taylor, J., Burke, J. & Osborne, J. (2003) Pilot evaluation of summer flounder stock enhancement potential using experimental ecology. Marine Ecology Progress Series 250, 263–278.

King, R.B. & Stanford, K.M. (2006) Headstarting as a management tool: A case study of the plains gartersnake. Herpetologica 62, 282–292.

Kingsbury, B.A. & Attum, O. (2009) Conservation strategies: captive rearing, translocation, and repatriation. In Snakes: Ecology and Conservation (eds S.J. Mullin & Seigel), pp. 202–220. Cornell University Press.

Klütsch, C.F.C., Maduna, S.N., Polikarpova, N., Forfang, K., Aspholm, P.E., Nyman, T., Eiken, H.G., Amundsen, P.A. & Hagen, S.B. (2019) Genetic changes caused by restocking and hydroelectric dams in demographically bottlenecked brown trout in a transnational subarctic riverine system. Ecology and Evolution 9, 6068–6081.

Knox, C.D., Jarvie, S., Easton, L.J. & Monks, J.M. (2017) Soft-Release, but not cool winter temperatures, reduces post-translocation dispersal of jewelled geckos. Journal of Herpetology 51, 490–496.

Knudsen, C.M., Shroder, S.L., Busack, C., Johnston, M. V., Pearsons, T.N. & Strom, C.R. (2008) Comparison of female reproductive traits and progeny of first-generation hatchery and wild upper Yakima River spring Chinook salmon. Transactions of the American Fisheries Society 137, 1433–1445.

Kock, T.J., Perry, R.W., Pope, A.C., Serl, J.D., Kohn, M. & Liedtke, T.L. (2018) Responses of hatchery- and natural-origin adult spring Chinook salmon to a trap-and-haul reintroduction program. North American Journal of Fisheries Management 38, 1004–1016.

Kołodziej-Sobocińska, M., Demiaszkiewicz, A.W., Pyziel, A.M. & Kowalczyk, R. (2018) Increased parasitic load in captive-released European bison (*Bison bonasus*) has important implications for reintroduction programs. EcoHealth 15, 467–471.

Koricheva, J., Jennions, M.D. & Lau, J. (2013) Temporal trends in effect sizes: causes, detection, and implications. In Handbook of meta-analysis in ecology and evolution (eds J. Koricheva, J. Gurevitch & K. Mengersen), pp. 237–254. Princeton University Press.

Kozłowski, J. (1993) Measuring fitness in life-history studies. Trends in Ecology and Evolution 8, 84–85.

Kraaijeveld-Smit, F.J.L., Griffiths, R.A., Moore, R.D. & Beebee, T.J.C. (2006) Captive breeding and the fitness of reintroduced species: a test of the responses to predators in a threatened amphibian. Journal of Applied Ecology 43, 360–365.

Kraus, B.T., Mccallen, E.B. & Williams, R.N. (2017) Evaluating the survival of translocated adult and captive-reared, juvenile eastern hellbenders (*Cryptobranchus alleganiensis alleganiensis*). Herpetologica 73, 271–276.

Krochmal, A.R., Roth, T.C. & O’malley, H. (2018) An empirical test of the role of learning in translocation. Animal Conservation 21, 36–44.

Laikre, L., Schwartz, M.K., Waples, R.S. & Ryman, N. (2010) Compromising genetic diversity in the wild: unmonitored large-scale release of plants and animals. Trends in Ecology and Evolution 25, 520–529.

Lajeunesse, M.J., Rosenberg, M.S. & Jennions, M. (2013) Phylogenetic nonindependence and meta-analysis. In Handbook of meta-analysis in ecology and evolution (eds J. Kuricheva, J. Gurevitch & K. Mengerson), pp. 284–299. Princeton University Press.

Lande, R. (1998) Anthropogenic, ecological and genetic factors in extinction and conservation. Population Ecology 40, 259–269.

Lapidge, S.J. (2005) Reintroduction increased vitamin E and condition in captive-bred yellow-footed rock wallabies *Petrogale xanthopus*. Oryx 39, 56–64.

Larsen, D.A., Harstad, D.L., Strom, C.R., Johnston, M. V., Knudsen, C.M., Fast, D.E., Pearsons, T.N. & Beckman, B.R. (2013) Early life history variation in hatchery- and natural-origin spring Chinook salmon in the Yakima River, Washington. Transactions of the American Fisheries Society 142, 540–555.

Larsson, S., Linnansaari, T., Vatanen, S., Serrano, I. & Haikonen, A. (2011) Feeding of wild and hatchery reared Atlantic salmon (Salmo salar L.) smolts during downstream migration. Environmental Biology of Fishes 92, 361–369.

Larsson, S., Serrano, I., Eriksson, L.-O. & Fleming, I. (2012) Effects of muscle lipid concentration on wild and hatchery brown trout (*Salmo trutta*) smolt migration. Canadian Journal of Fisheries & Aquatic Sciences 69, 1–12.

Lebata, Ma.J.H.L., LE Vay, L., Walton, M.E., Biñas, J.B., Quinitio, E.T., Rodriguez, E.M. & Primavera, J.H. (2009) Evaluation of hatchery-based enhancement of the mud crab, *Scylla spp*., fisheries in mangroves: comparison of species and release strategies. Marine and Freshwater Research 60, 58.

Lebata-Ramos, Ma.J.H., Doyola-Solis, E.F.C., Abrogueña, J.B.R., Ogata, H., Sumbing, J.G. & Sibonga, R.C. (2013) Evaluation of post-release behavior, recapture, and growth rates of hatchery-reared abalone *Haliotis asinina* released in Sagay Marine Reserve, Philippines. Reviews in Fisheries Science 21, 433–440.

Lee, J.-H. & Park, D.-S. (2011) Spatial ecology of translocated and resident Amur ratsnakes (*Elaphe schrenckii*) in two mountain valleys of South Korea. Asian Herpetological Research 2, 223–229.

Lehrer, E.W., Schooley, R.L., Nevis, J.M., Kilgour, R.J., Wolff, P.J. & Magle, S.B. (2016) Happily ever after? Fates of translocated nuisance woodchucks in the Chicago metropolitan area. Urban Ecosystems 19, 1389–1403.

Lenormand, T. (2002) Gene flow and the limits to natural selection. Trends in Ecology & Evolution 17, 183–189.

Lenth, R. V. (2021) emmeans: Estimated marginal means, aka least-square means.

Lepeigneul, O., Ballouard, J.M., Bonnet, X., Beck, E., Barbier, M., Ekori, A., Buisson, E. & Caron, S. (2014) Immediate response to translocation without acclimation from captivity to the wild in Hermann’s tortoise. European Journal of Wildlife Research 60, 897– 907.

Lettink, M. (2007) Detectability, movements and apparent lack of homing in *Hoplodactylus maculatus* (Reptilia: Diplodactylidae) following translocation. New Zealand Journal of Ecology 31, 111–116.

Linklater, W.L., Adcock, K., Du Preez, P., Swaisgood, R.R., Law, P.R., Knight, M.H., Gedir, J. V. & Kerley, G.I.H. (2011) Guidelines for large herbivore translocation simplified: black rhinoceros case study. Journal of Applied Ecology 48, 493–502.

Lino, P.G., Bentes, L., Abecasis, D., Santos, M.N.D. & Erzini, K. (2009) Comparative behavior of wild and hatchery reared white sea bream (*Diplodus sargus*) released on artificial reefs off the Algarve (Southern Portugal). In Tagging and Tracking of Marine Animals with Electronic Devices (eds J.L. Nielsen, H. Arrizabalaga, N. Fragoso, A. Hobday, M. Lutcavage & J. Sibert), pp. 23–34. Springer Netherlands, Dordrecht.

Lumsden, H.G. & Drever, M.C. (2002) Overview of the trumpeter swan reintroduction program in Ontario, 1982-2000. Waterbirds: The International Journal of Waterbird Biology 25, 301–312.

Lutz, M.L., Tonkin, Z., Yen, J.D.L., Johnson, G., Ingram, B.A., Sharley, J., Lyon, J., Chapple, D.G., Sunnucks, P. & Pavlova, A. (2021) Using multiple sources during reintroduction of a locally extinct population benefits survival and reproduction of an endangered freshwater fish. Evolutionary Applications 14, 950–964.

Lynch, M. (1991) Methods for the analysis of comparative data in evolutionary biology. Evolution 45, 1065–1080.

Lynch, M. & O’hely, M. (2001) Captive breeding and the genetic fitness of natural populations. Conservation Genetics 2, 363–378.

Mann, K.A., Marty Holtgren, J., Auer, N.A. & Ogren, S.A. (2011) Comparing size, movement, and habitat selection of wild and streamside-reared lake sturgeon. North American Journal of Fisheries Management 31, 305–314.

Mathews, F., Orros, M., Mclaren, G., Gelling, M. & Foster, R. (2005) Keeping fit on the ark: assessing the suitability of captive-bred animals for release. Biological Conservation 121, 569–577.

Mccallen, E.B., Kraus, B.T., Burgmeier, N.G., Fei, S. & Williams, R.N. (2018) Movement and habitat use of eastern hellbenders (*Cryptobranchus alleganiensis alleganiensis*) following population augmentation. Herpetologica 74, 283–293.

Mccleery, R., Oli, M.K., Hostetler, J.A., Karmacharya, B., Greene, D., Winchester, C., Gore, J., Sneckenberger, S., Castleberry, S.B. & Mengak, M.T. (2013) Are declines of an endangered mammal predation-driven, and can a captive-breeding and release program aid their recovery? Journal of Zoology 291, 59–68.

Melnychuk, M.C., Korman, J., Hausch, S., Welch, D.W., Mccubbing, D.J.F. & Walters, C.J. (2014) Marine survival difference between wild and hatchery-reared steelhead trout determined during early downstream migration. Canadian Journal of Fisheries and Aquatic Sciences 71, 831–846.

Melstrom, R.T., Salau, K.R. & Shanafelt, D.W. (2019) The optimal timing of reintroducing captive populations into the wild. Ecological Economics 156, 174–184.

Mengersen, K., Schmid, C.H., Jennions, M.D., & Gurevitch, J. (2013) Statistical models and approaches to inference. In Handbook of meta-analysis in ecology and evolution (eds J. Koricheva, J. Gurevitch & K. Mengersen), pp. 89-107. Princeton University Press.

Miller, K.A., Bell, T.P., & Germano, J.M. (2014) Understanding publication bias in reintroduction biology by assessing translocations of New Zealand’s herpetofauna. Conservation Biology 28, 1045–1056.

Mills, D.J., Gardner, C. & Johnson, C.R. (2006) Experimental reseeding of juvenile spiny lobsters (*Jasus edwardsii*): Comparing survival and movement of wild and naïve lobsters at multiple sites. Aquaculture 254, 256–268.

Moher, D., Liberati, A., Tetzlaff, J., Altman, D.G., Altman, D., Antes, G., Atkins, D., Barbour, V., Barrowman, N., Berlin, J.A., Clark, J., Clarke, M., Cook, D., D’amico, R., Deeks, J.J., et al. (2009) Preferred reporting items for systematic reviews and meta-analyses: The PRISMA statement. PLoS Medicine 6.

Molony, S.E., Dowding, C. V., Baker, P.J., Cuthill, I.C. & Harris, S. (2006) The effect of translocation and temporary captivity on wildlife rehabilitation success: An experimental study using European hedgehogs (*Erinaceus europaeus*). Biological Conservation 130, 530–537.

Monzyk, F.R., Jonasson, B.C., Hoffnagle, T.L., Keniry, P.J., Carmichael, R.W. & Cleary, P.J. (2009) Migration characteristics of hatchery and natural spring Chinook salmon smolts from the Grande Ronde River Basin, Oregon, to Lower Granite Dam on the Snake River. Transactions of the American Fisheries Society 138, 1093–1108.

Moore, M., Berejikian, B.A. & Tezak, E.P. (2012) Variation in the early marine survival and behavior of natural and hatchery-reared Hood Canal steelhead. PLoS ONE 7, e49645.

Moore, S.J. & Battley, P.F. (2006) Differences in the digestive organ morphology of captive and wild brown teal *Anas chlorotis* and implications for releases. Bird Conservation International 16, 253–264.

Moritz, C. (2002) Strategies to protect biological diversity and the evolutionary processes that sustain it. Systematic Biology 51, 238–254.

Morris, S.D., Brook, B.W., Moseby, K.E., & Johnson, C.N. (2021) Factors affecting success of conservation translocations of terrestrial vertebrates: a global systematic review. Global Ecology and Conservation 28, e01630.

Moseby, K.E., Hill, B.M. & Lavery, T.H. (2014) Tailoring release protocols to individual species and sites: one size does not fit all. PLoS ONE 9.

Moyes, K., Morgan, B.J.T., Morris, A., Morris, S.J., Clutton-Brock, T.H. & Coulson, T. (2009) Exploring individual quality in a wild population of red deer. Journal of Animal Ecology 78, 406–413.

Mulder, K.P., Walde, A.D., Boarman, W.I., Woodman, A.P., Latch, E.K. & Fleischer, R.C. (2017) No paternal genetic integration in desert tortoises (*Gopherus agassizii*) following translocation into an existing population. Biological Conservation 210, 318–324.

Nakagawa, S. & Santos, E.S.A. (2012) Methodological issues and advances in biological meta-analysis. Evolutionary Ecology 26, 1253–1274.

Nakagawa, S. & Cuthill, I. C. (2007) Effect size, confidence interval and statistical significant: a practical guide for biologists. Biological Reviews 82: 591–605.

Nash, D.J. & Griffiths, R.A. (2018) Ranging behaviour of adders (*Vipera berus*) translocated from a development site. Herpetological Journal 28, 155–159.

Nason, S.E., Lloyd, N., Kelly, C.D., Brichieri-Colombi, T.A., Dalrymple, S.E. & Moehrenschlager, A. (2021) Maximizing the effectiveness of qualitative systematic reviews: A case study on terrestrial arthropod conservation translocations. Biological Conservation 254, 108948.

Nazar, F.N. & Marin, R.H. (2011) Chronic stress and environmental enrichment as opposite factors affecting the immune response in Japanese quail (*Coturnix coturnix japonica*). Stress 14, 166–173.

Nelson, T.C., Rosenau, M.L. & Johnston, N.T. (2005) Behavior and survival of wild and hatchery-origin winter steelhead spawners caught and released in a recreational fishery. North American Journal of Fisheries Management 25, 931–943.

Neuman, K., Stenzel, L., Warriner, J., Page, G., Erbes, J., Eyster, C., Miller, E. & Henkel, L. (2013) Success of captive-rearing for a threatened shorebird. Endangered Species Research 22, 85–94.

Newmark, W.D., Jenkins, C.N., Pimm, S.L., Mcneally, P.B. & Halley, J.M. (2017) Targeted habitat restoration can reduce extinction rates in fragmented forests. Proceedings of the National Academy of Sciences of the United States of America 114, 9635–9640.

Nicoll, M.A.C., Jones, C.G. & Norris, K. (2004) Comparison of survival rates of captive-reared and wild-bred Mauritius kestrels (*Falco punctatus*) in a re-introduced population. Biological Conservation 118, 539–548.

Noble, D.W.A., Lagisz, M., O’dea, R.E. & Nakagawa, S. (2017) Nonindependence and sensitivity analyses in ecological and evolutionary meta-analyses. Molecular Ecology 26, 2410–2425.

van Noordwijk, A.J. & De Jong, G. (1986) Acquisition and allocation of resources: their influence on variation in life history tactics. The American Naturalist 128, 137–142.

Norris, T.A., Littnan, C.L. & Gulland, F.M.D. (2011) Evaluation of the captive care and post-release behavior and survival of seven juvenile female Hawaiian monk seals (*Monachus schauinslandi*). Aquatic Mammals 37, 342–353.

Nussear, K.E., Tracy, C.R., Medica, P.A., Wilson, D.S., Marlow, R.W. & Corn, P.S. (2012) Translocation as a conservation tool for Agassiz’s desert tortoises: survivorship, reproduction, and movements. The Journal of Wildlife Management 76, 1341–1353.

Nyqvist, D., Mccormick, S.D., Greenberg, L., Ardren, W.R., Bergman, E., Calles, O. & Castro-Santos, T. (2017) Downstream migration and multiple dam passage by Atlantic salmon smolts. North American Journal of Fisheries Management 37, 816–828.

O’dea, R.E., Lagisz, M., Jennions, M.D., Koricheva, J., Noble, D.W.A., Parker, T.H., Gurevitch, J., Page, M.J., Stewart, G., Moher, D. & Nakagawa, S. (2021) Preferred reporting items for systematic reviews and meta-analyses in ecology and evolutionary biology: a PRISMA extension. Biological Reviews 96, 1695–1722.

Økland, F., Heggberget, T.G. & Jonsson, B. (1995) Migratory behaviour of wild and farmed Atlantic salmon (*Salmo salar*) during spawning. Journal of Fish Biology 46, 1–7.

Olivier, J., & Bell, M. L. (2013) Effect sizes for 2 × 2 contingency tables. PSoS ONE 8, e5877.

Osterback, A.-M.K., Frechette, D.M., Hayes, S.A., Bond, M.H., Shaffer, S.A. & Moore, J.W. (2014) Linking individual size and wild and hatchery ancestry to survival and predation risk of threatened steelhead (*Oncorhynchus mykiss*). Canadian Journal of Fisheries and Aquatic Sciences 71, 1877–1887.

Ostermann, S.D., Deforge, J.R. & Edge, W.D. (2001) Captive breeding and reintroduction evaluation criteria: a case study of peninsular bighorn sheep. Conservation Biology 15, 749– 760.

Parish, D.M.B. & Sotherton, N.W. (2007) The fate of released captive-reared grey partridges *Perdix perdix*: implications for reintroduction programmes. Wildlife Biology 13, 140–149.

Parker, G. (1979) Sexual selection and sexual conflict. In Sexual Selection and Reproductive Competition in Insects (ed M. Blum), pp. 123–166. New York Academic Press.

Peck, A.J., Harris, J.L., Farris, J.L. & Christian, A.D. (2014) Survival and horizontal movement of the freshwater mussel *Potamilus capax* (Green, 1832) following relocation within a Mississippi Delta stream system. American Midland Naturalist 172, 76–90.

Pedrono, M. & Sarovy, A. (2000) Trial release of the world’s rarest tortoise *Geochelone yniphora* in Madagascar. Biological Conservation 95, 333–342.

Pekkala, N., Kotiaho, J.S. & Puurtinen, M. (2011) Laboratory relationships between adult lifetime reproductive success and fitness surrogates in a *Drosophila littoralis* population. PLoS ONE 6, e24560.

Pérez, I., Anadón, J.D., Díaz, M., Nicola, G.G., Tella, J.L. & Giménez, A. (2012) What is wrong with current translocations? A review and a decision-making proposal. Frontiers in Ecology and the Environment 10, 494–501.

Pérez-Buitrago, N., García, M., Sabat, A., Delgado, J., Álvarez, A., Mcmillan, O. & Funk, S.. (2008) Do headstart programs work? Survival and body condition in headstarted Mona Island iguanas Cyclura cornuta stejnegeri. Endangered Species Research 6, 55–65.

Piggins, D.J. & Mills, C.P.R. (1985) Comparative aspects of the biology of naturally produced and hatchery-reared Atlantic salmon smolts (*Salmo salar L*.). Aquaculture 45, 321–333.

Pille, F., Caron, S., Bonnet, X., Deleuze, S., Busson, D., Etien, T., Girard, F. & Ballouard, J.-M. (2018) Settlement pattern of tortoises translocated into the wild: a key to evaluate population reinforcement success. Biodiversity and Conservation 27, 437–457.

Pimm, S.L., Jenkins, C.N., Abell, R., Brooks, T.M., Gittleman, J.L., Joppa, L.N., Raven, P.H., Roberts, C.M. & Sexton, J.O. (2014) The biodiversity of species and their rates of extinction, distribution, and protection. Science 344, 987.

Pinter, K., Epifanio, J. & Unfer, G. (2019) Release of hatchery-reared brown trout (*Salmo trutta*) as a threat to wild populations? A case study from Austria. Fisheries Research 219, 105296.

Plummer, M.V. & Mills, N.E. (2000) Spatial ecology and survivorship of resident and translocated hognose snakes (*Heterodon platirhinos*). Journal of Herpetology 34, 565–575.

Poirier, M.-A. & Festa-Bianchet, M. (2018) Social integration and acclimation of translocated bighorn sheep (*Ovis canadensis*). Biological Conservation 218, 1–9.

Poisot, T. (2011) The digitize package: extracting numerical data from scatterplots. The R Journal 3, 26–27.

Poo, S., Bogisich, A., Mack, M., Lynn, B.K. & Devan-Song, A. (2022) Post-release comparisons of amphibian growth reveal challenges with sperm cryopreservation as a conservation tool. Conservation Science and Practice 4, 1–9.

Poulin, R.G., Todd, L.D., Wellicome, T.I. & Brigham, R.M. (2006) Assessing the feasibility of release techniques for captive-bred burrowing owls. Journal of Raptor Research 40, 142– 150.

Powell, M.S., Hardy, R.W., Flagg, T.A. & Kline, P.A. (2010) Proximate composition and fatty acid differences in hatchery-reared and wild Snake River Sockeye salmon overwintering in nursery lakes. North American Journal of Fisheries Management 30, 530– 537.

, A.S. & Knight, T.M. (2001) Effectiveness in conservation practice: pointers from medicine and public health. Conservation Biology 15, 50–54.

Quinn, T.P., Seamons, T.R. & Johnson, S.P. (2012) Stable isotopes of carbon and nitrogen indicate differences in marine ecology between wild and hatchery-produced steelhead. Transactions of the American Fisheries Society 141, 526–532.

Rantanen, E.M., Buner, F., Riordan, P., Sotherton, N. & Macdonald, D.W. (2010) Vigilance, time budgets and predation risk in reintroduced captive-bred grey partridges Perdix perdix. Applied Animal Behaviour Science 127, 43–50.

Ray, M., Stoner, A. & O’connell, S. (1994) Size-specific predation of juvenile queen conch *Strombus gigas*: implications for stock enhancement. Aquaculture 128, 79–88.

Reading, R.P., Clark, T.W., & Griffith, B. (1997) The influence of valuational and organization consideration on the succes of rare species translocations. Biological Conservation 79, 217–225.

Reading, R.P., Miller, B. & Shepherdson, D. (2013) The value of enrichment to reintroduction success. Zoo Biology 32, 332–341.

Reid, J.M., Arcese, P., Nietlisbach, P., Wolak, M.E., Muff, S., Dickel, L., & Keller, L.F. (2021) Immigration counter-acts local micro-evolution of a major fitness component: migration-selection balance in free-living song sparrows. Evolution Letters 5, 48–60.

Reinert, H.K. & Rupert, R.R. (1999) Impacts of translocation on behavior and survival of timber rattlesnakes, *Crotalus horridus*. Journal of Herpetology 33, 45–61.

Resende, P.S., Viana-Junior, A.B., Young, R.J. & Azevedo, C.S. (2021) What is better for animal conservation translocation programmes: soft-or hard-release? A phylogenetic meta-analytical approach. Journal of Applied Ecology 58, 1122–1132.

Rittenhouse, C.D., Millspaugh, J.J., Hubbard, M.W. & Sheriff, S.L. (2007) Movements of translocated and resident three-toed box turtles. Journal of Herpetology 41, 115–121.

Roche, E.A., Cuthbert, F.J. & Arnold, T.W. (2008) Relative fitness of wild and captive-reared piping plovers: does egg salvage contribute to recovery of the endangered Great Lakes population? Biological Conservation 141, 3079–3088.

Roe, J.H., Frank, M.R. & Kingsbury, B.A. (2015) Experimental evaluation of captive-rearing practices to improve success of snake reintroductions. Herpetological Conservation and Biology 10, 711–722.

Roff, D.A. (2002) *Life History Evolution*. Sinauer Associates, Inc. Sunderland, Ma, USA.

Rosenberg, M.S., Rothstein, H.R. & Gurevitch, J. (2013) Effect sizes: Conventional choices and calculations. In Handbook of meta-analysis in ecology and evolution (eds J. Kuricheva, J. Gurevitch & K. Mengersen), pp. 61–71. Princeton University Press.

Rosenthal, R. & Rubin, D.B. (1982) Comparing effect sizes of independent studies. Psychological Bulletin 92, 500–504.

Ruckstuhl, K.E. & Neuhaus, P. (2005) Sexual Segregation in Vertebrates: Ecology of the two sexes. Cambridge University Press.

Rummel, L., Martínez-Abraín, A., Mayol, J., Ruiz-Olmo, J., Mañas, F., Jiménez, J., Gómez, J.A. & Oro, D. (2016) Use of wild-caught individuals as a key factor for success in vertebrate translocations. Animal Biodiversity and Conservation 39, 207–219.

Ruth, T.K., Logan, K.A., Sweanor, L.L., Hornocker, M.G. & Temple, L.J. (1998) Evaluating Cougar Translocation in New Mexico. The Journal of Wildlife Management 62, 1264–1275.

Ryan, B.A., Smith, S.G., Butzerin, J.A.M. & Ferguson, J.W. (2003) Relative vulnerability to avian predation of juvenile salmonids tagged with passive integrated transponders in the Columbia River estuary, 1998–2000. Transactions of the American Fisheries Society 132, 275–288.

Ryman, N. & Laikre, L. (1991) Effects of supportive breeding on the genetically effective population size. Conservation Biology 5, 325–329.

Sah, P., Nussear, K.E., Esque, T.C., Aiello, C.M., Hudson, P.J. & Bansal, S. (2016) Inferring social structure and its drivers from refuge use in the desert tortoise, a relatively solitary species. Behavioral Ecology and Sociobiology 70, 1277–1289.

Saloniemi, I., Jokikokko, E., Kallio-Nyberg, I., Jutila, E. & Pasanen, P. (2004) Survival of reared and wild Atlantic salmon smolts: size matters more in bad years. ICES Journal of Marine Science 61, 782–787.

Santos, T., Pérez-Tris, J., Carbonell, R., Tellería, J.L. & Díaz, J.A. (2009) Monitoring the performance of wild-born and introduced lizards in a fragmented landscape: implications for ex situ conservation programmes. Biological Conservation 142, 2923–2930.

Satake, A. & Araki, H. (2012) Stocking of captive-bred fish can cause long-term population decline and gene pool replacement: predictions from a population dynamics model incorporating density-dependent mortality. Theoretical Ecology 5, 283–296.

Scheele, B.C., Hollanders, M., Hoffmann, E.P., Newell, D.A., Lindenmayer, D.B., Mcfadden, M., Gilbert, D.J. & Grogan, L.F. (2021) Conservation translocations for amphibian species threatened by chytrid fungus: a review, conceptual framework, and recommendations. Conservation Science and Practice 3, 1–15.

Scillitani, L., Sturaro, E., Menzano, A., Rossi, L., Viale, C. & Ramanzin, M. (2012) Post-release spatial and social behaviour of translocated male Alpine ibexes (*Capra ibex ibex*) in the eastern Italian Alps. European Journal of Wildlife Research 58, 461–472.

Seddon, P.J., Armstrong, D.P. & Maloney, R.F. (2007) Developing the science of reintroduction biology. Conservation Biology 21, 303–312.

Seddon, P.J., Griffiths, C.J., Soorae, P.S. & Armstrong, D.P. (2014) Reversing defaunation: Restoring species in a changing world. Science 345, 406–411.

Senior, A.M., Grueber, C.E., Kamiya, T., Lagisz, M., O’dwyer, K., Santos, E.S.A. & Nakagawa, S. (2016) Heterogeneity in ecological and evolutionary meta-analyses: its magnitude and implications. Ecology 97, 3293–3299.

Serrano, I., Larsson, S. & Eriksson, L.-O. (2009) Migration performance of wild and hatchery sea trout (*Salmo trutta L.*) smolts—implications for compensatory hatchery programs. Fisheries Research 99, 210–215.

Shaw, R.G. (2019) From the past to the future: considering the value and limits of evolutionary Prediction. The American Naturalist 193, 1–10.

Shier, D.M. & Owings, D.H. (2007) Effects of social learning on predator training and postrelease survival in juvenile black-tailed prairie dogs, *Cynomys ludovicianus*. Animal Behaviour 73, 567–577.

Sinn, D.L., Cawthen, L., Jones, S.M., Pukk, C. & Jones, M.E. (2014) Boldness towards novelty and translocation success in captive-raised, orphaned Tasmanian devils. Zoo Biology 33, 36–48.

Skaala, O., Jørstad, K. & Borgstrøm, R. (2011) Genetic impact on two wild brown trout (*Salmo trutta*) populations after release of non-indigenous hatchery spawners. Canadian Journal of Fisheries and Aquatic Sciences 53, 2027–2035.

Skikne, S.A., Borker, A.L., Terrill, R.S. & Zavaleta, E. (2020) Predictors of past avian translocation outcomes inform feasibility of future efforts under climate change. Biological Conservation 247, 108597.

Slaugh, B.T., Flinders, J.T., Roberson, J.A. & Johnston, N.P. (1992) Effect of rearing method on chukar survival. The Great Basin Naturalist 52, 25–28.

Snyder, N.F.R., Derrickson, S.R., Beissinger, S.R., Wiley, J.W., Smith, T.B., Toone, W.D. & Miller, B. (1996) Limitations of captive breeding in endangered species recovery. Conservation Biology 10, 338–348.

Stamps, J.A. & Swaisgood, R.R. (2007) Someplace like home: experience, habitat selection and conservation biology. Applied Animal Behaviour Science 102, 392–409.

Stark, E.J., Vidergar, D.T., Kozfkay, C.C. & Kline, P.A. (2018) Egg viability and egg-to-fry survival of captive-reared Chinook salmon released to spawn naturally. Transactions of the American Fisheries Society 147, 128–138.

Stewart, D.C. (2015) Ranching to the rod: an evaluation of adult returns from hatchery-reared Atlantic salmon smolts released in Scottish rivers. Scottish Marine and Freshwater Science 6, 1–13.

Stewart, G. (2010) Meta-analysis in applied ecology. Biology Letters 6, 78–81.

Stoner, A. & Davis, M. (1994) Experimental outplanting of juvenile queen conch, Strombus gigas: comparison of wild and hatchery-reared stocks. Fishery Bulletin-National Oceanic and Atmospheric Administration 92, 390–411.

Stringwell, R., Lock, A., Stutchbury, C.J., Baggett, E., Taylor, J., Gough, P.J. & Leaniz, C.G. DE (2014) Maladaptation and phenotypic mismatch in hatchery-reared Atlantic salmon *Salmo salar* released in the wild. Journal of Fish Biology 85, 1927–1945.

Stuparyk, B., Horn, C.J., Karabatsos, S. & Arteaga, J. (2018) A meta-analysis of animal survival following translocation: comparisons between conflict and conservation efforts. Canadian Wildlife Biology and Managment 7, 3–17.

Sturdevant, M.V., Fergusson, E., Hillgruber, N., Reese, C., Orsi, J., Focht, R., Wertheimer, A. & Smoker, B. (2012) Lack of trophic competition among wild and hatchery juvenile chum salmon during early marine residence in Taku Inlet, Southeast Alaska. Environmental Biology of Fishes 94, 101–116.

Sulak, K.J., Randall, M.T. & Clugston, J.P. (2014) Survival of hatchery Gulf sturgeon (Acipenser oxyrinchus desotoi Mitchill, 1815) in the Suwannee River, Florida: a 19-year evaluation. Journal of Applied Ichthyology 30, 1428–1440.

Sullivan, B.K., Nowak, E.M. & Kwiatkowski, M.A. (2015) Problems with mitigation translocation of herpetofauna. Conservation Biology 29, 12–18.

Sutherland, W.J., Pullin, A.S., Dolman, P.M. & Knight, T.M. (2004) The need for evidence-based conservation. Trends in Ecology and Evolution 19, 305–308.

Tarka, M., Guenther, A., Niemelä, P.T., Nakagawa, S. & Noble, D.W.A. (2018) Sex differences in life history, behavior, and physiology along a slow-fast continuum: a meta-analysis. Behavioral Ecology and Sociobiology 72. Behavioral Ecology and Sociobiology.

Taylor, M.D., Laffan, S.W., Fairfax, A.V. & Payne, N.L. (2017) Finding their way in the world: Using acoustic telemetry to evaluate relative movement patterns of hatchery-reared fish in the period following release. Fisheries Research 186, 538–543.

Teixeira, C.P., De Azevedo, C.S., Mendl, M., Cipreste, C.F. & Young, R.J. (2007) Revisiting translocation and reintroduction programmes: the importance of considering stress. Animal Behaviour 73, 1–13.

Temsiripong, Y., Woodward, A.R., Ross, J.P., Kubilis, P.S. & Percival, H.F. (2006) Survival and growth of american alligator (*Alligator mississippiensis*) hatchlings after artificial incubation and repatriation. Journal of Herpetology 40, 415–423.

Terhune, T.M., Sisson, D.C. & Stribling, H.L. (2006) The efficacy of relocating wild northern bobwhites prior to breeding season. The Journal of Wildlife Management 70, 914– 921.

Tetzlaff, S.J., Sperry, J.H. & Degregorio, B.A. (2018) Captive-reared juvenile box turtles innately prefer naturalistic habitat: Implications for translocation. Applied Animal Behaviour Science 204, 128–133.

Tetzlaff, S.J., Sperry, J.H. & Degregorio, B.A. (2019a) Effects of antipredator training, environmental enrichment, and soft release on wildlife translocations: a review and meta-analysis. Biological Conservation 236, 324–331.

Tetzlaff, S.J., Sperry, J.H., Kingsbury, B.A. & Degregorio, B.A. (2019b) Captive-rearing duration may be more important than environmental enrichment for enhancing turtle head-starting success. Global Ecology and Conservation 20, e00797.

Thoday, J.M. (1953) Components of fitness. Symposium of the Society for Experimental Biology 8, 96–113.

Thompson, B.C., Porak, W.F., Leone, E.H. & Allen, M.S. (2016) Using radiotelemetry to compare the initial behavior and mortality of hatchery-reared and wild juvenile Florida bass. Transactions of the American Fisheries Society 145, 374–385.

Thorstad, E.B., Heggberget, T.G. & Økland, F. (1998) Migratory behaviour of adult wild and escaped farmed Atlantic salmon, *Salmo salar L.*, before, during and after spawning in a Norwegian river. Aquaculture Research 29, 419–428.

Thorstad, E.B., Økland, F., Finstad, B., Sivertsgård, R., Plantalech, N., Bjørn, P.A. & Mckinley, R.S. (2007) Fjord migration and survival of wild and hatchery-reared Atlantic salmon and wild brown trout post-smolts. Hydrobiologia 582, 99–107.

Trested, D.G., Ware, K., Bakal, R. & Isely, J.J. (2011) Microhabitat use and seasonal movements of hatchery-reared and wild shortnose sturgeon in the Savannah River, South Carolina – Georgia. Journal of Applied Ichthyology 27, 454–461.

Trivers, R.L. (1972) Parental investment and sexual selection. In Sexual Selection and the Descent of Man, 1871-1971 pp. 136–179. Aldine, Chicago, IL.

Tsakiris, E.T., Randklev, C.R., Blair, A., Fisher, M. & Conway, K.W. (2017) Effects of translocation on survival and growth of freshwater mussels within a West Gulf Coastal Plain river system. Aquatic Conservation: Marine and Freshwater Ecosystems 27, 1240–1250.

Tuberville, T.D., Clark, E.E., Buhlmann, K.A. & Gibbons, J.W. (2005) Translocation as a conservation tool: Site fidelity and movement of repatriated gopher tortoises (*Gopherus polyphemus*). Animal Conservation 8, 349–358.

Turek, J., Horký, P., Velíšek, J., Slavík, O., Hanák, R. & Randák, T. (2010a) Recapture rate and growth of hatchery-reared brown trout (*Salmo trutta v. fario, L*.) in Blanice River and the effect of stocking on wild brown trout and grayling (*Thymallus thymallus*, L.). Journal of Applied Ichthyology 26, 881–885.

Turek, J., Randák, T., Horký, P., Zllábek, V., Velíšek, J., Slavík, O. & Hanák, R. (2010b) Post-release growth and dispersal of pond and hatchery-reared European grayling Thymallus thymallus compared with their wild conspecifics in a small stream. Journal of Fish Biology 76, 684–693.

Turlure, C., Radchuk, V., Baguette, M., Meijrink, M., van den Burg, A., De Vries, M.W. & van Duinen, G.J. (2013) Plant quality and local adaptation undermine relocation in a bog specialist butterfly. Ecology and Evolution 3, 244–254.

Tweed, E.J., Foster, J.T., Woodworth, B.L., Monahan, W.B., Kellerman, J.L. & Lieberman, A. (2006) Breeding biology and success of a reintroduced population of the critically endangered Puaiohi (*Myadestes Palmeri*). The Auk 123, 753–763.

Unger, C. & Klaus, Siegfried (2009) Lebenserwartung und Verlustursachen umgesiedelter Auerhühner *Tetrao urogallus* in Thüringen. Ornithologischer Anzeiger 48, 50–55.

Urke, H.A., Kristensen, T., Ulvund, J.B. & Alfredsen, J.A. (2013) Riverine and fjord migration of wild and hatchery-reared Atlantic salmon smolts. Fisheries Management and Ecology 20, 544–552.

Vanderwerf, E.A., Crampton, L.H., Diegmann, J.S., Atkinson, C.T. & Leonard, D.L. (2014) Survival estimates of wild and captive-bred released Puaiohi, an endangered Hawaiian thrush. The Condor: Ornithological Applications 116, 609–618.

Viechtbauer, W. (2010) Conducting meta-analyses in R with the metafor. Journal of Statistical Software 36, 1–48.

Viechtbauer, W. & Cheung, M.W.-L. (2010) Outlier and influence diagnostics for meta-analysis. Research Synthesis Methods 1, 112–125.

Wake, D.B. & Vredenburg, V.T. (2008) Are we in the midst of the sixth mass extinction? A view from the world of amphibians. Proceedings of the National Academy of Sciences of the United States of America 105, 11466–11473.

Waples, R.S., Hindar, K., Karlsson, S. & Hard, J.J. (2016) Evaluating the Ryman–Laikre effect for marine stock enhancement and aquaculture. Current Zoology 62, 617–627.

Weigel, D.E., Connolly, P.J. & Powell, M.S. (2014) Fluvial rainbow trout contribute to the colonization of steelhead (*Oncorhynchus mykiss*) in a small stream. Environmental Biology of Fishes 97, 1149–1159.

Weise, F.J., Lemeris, J., Stratford, K.J., van Vuuren, R.J., Munro, S.J., Crawford, S.J., Marker, L.L. & Stein, A.B. (2015) A home away from home: insights from successful leopard (*Panthera pardus*) translocations. Biodiversity and Conservation 24, 1755–1774.

Welch, D.W., Ward, B.R. & Batten, S.D. (2004) Early ocean survival and marine movements of hatchery and wild steelhead trout (*Oncorhynchus mykiss*) determined by an acoustic array: Queen Charlotte Strait, British Columbia. Deep Sea Research Part II: Topical Studies in Oceanography 51, 897–909.

Williamson, K.S., Murdoch, A.R., Pearsons, T.N., Ward, E.J. & Ford, M.J. (2010) Factors influencing the relative fitness of hatchery and wild spring Chinook salmon (*Oncorhynchus tshawytscha*) in the Wenatchee River, Washington, USA. Canadian Journal of Fisheries and Aquatic Sciences 67, 1840–1851.

Wilson, A.J. & Nussey, D.H. (2010) What is individual quality? An evolutionary perspective. Trends in Ecology and Evolution 25, 207–214.

Wisely, S.M., Howard, J., Williams, S.A., Bain, O., Santymire, R.M., Bardsley, K.D. & Williams, E.S. (2008) An unidentified filiarial species and its impact on fitness in wild population of the black-footed ferret (*Mustela nigripes*). Journal of Wildlife Diseases 44, 53–64.

Wolf, C.M., Griffith, B., Reed, C. & Temple, S.A. (1996) Avian and mammalian translocations: update and reanalysis of 1987 survey data. Conservation Biology 10, 1142– 1154.

Wolfe, A.K., Fleming, P.A., Bateman, P.W., Wolfe, A.K., Fleming, P.A. & Bateman, P.W. (2018) Impacts of translocation on a large urban-adapted venomous snake. Wildlife Research 45, 316–324.

Wood, C., Welch, D., Godbout, L. & Cameron, J. (2012) Marine migratory behavior of hatchery-reared anadromous and wild non-anadromous sockeye salmon revealed by acoustic tags. American Fisheries Society Symposium 76, 289–311.

WWF/ZSL (2020) The Living Planet Index database. Www.livingplanetindex.org.

Zhang, Z., Gao, L. & Zhang, X. (2022) Environmental enrichment increases aquatic animal welfare: A systematic review and meta-analysis. Reviews in Aquaculture 14, 1120–1135.

Zhu, J., Gosnell, J.S., Akallal, L. & Goltsman, M. (2022) Fear changes traits and increases survival: a metalJanalysis evaluating the efficacy of antipredator training in captivelJrearing programs. Restoration Ecology 31, 1–11.

