## Appendix S1 for "The fitness consequences of wildlife conservation translocations: a meta-analysis"

to

The fitness consequences of wildlife conservation translocations: a meta-analysis

Iwo P. Gross[1,*], Alan E. Wilson[2], & Matthew E. Wolak[1]

[1]Department of Biological Sciences, Auburn University, Auburn, AL 36849, USA

[2]School of Fisheries, Aquaculture, and Aquatic Sciences, Auburn University, Auburn, AL 36849, USA

O'Dea, R.E., Lagisz, M., Jennions, M.D., Koricheva, J., Noble, D.W., Parker, T.H., Gurevitch, J., Page, M.J., Stewart, G., Moher, D. and Nakagawa, S. (2021), Preferred reporting items for systematic reviews and meta-analyses in ecology and evolutionary biology: a PRISMA extension. Biol Rev. doi:10.1111/brv.12721

| Checklist item | Sub-item number | Sub-item | Reported by authors? | Notes |
|---|---|---|---|---|
| Title and abstract | 1.1 | Identify the review as a systematic review, meta-analysis, or both | Yes | |
| | 1.2 | Summarise the aims and scope of the review | Yes | |
| | 1.3 | Describe the data set | Yes | |
| | 1.4 | State the results of the primary outcome | Yes | |
| | 1.5 | State conclusions | Yes | |
| | 1.6 | State limitations | Yes | |
| Aims and questions | 2.1 | Provide a rationale for the review | Yes | |
| | 2.2 | Reference any previous reviews or meta-analyses on the topic | Yes | |
| | 2.3 | State the aims and scope of the review (including its generality) | Yes | |
| | 2.4 | State the primary questions the review addresses (e.g. which moderators were tested) | Yes | |
| | 2.5 | Describe whether effect sizes were derived from experimental and/or observational comparisons | Yes | |

| | | | | |
|---|---|---|---|---|
| Review registration | 3.1 | Register review aims, hypotheses (if applicable), and methods in a time-stamped and publicly accessible archive and provide a link to the registration in the methods section of the manuscript. Ideally registration occurs before the search, but it can be done at any stage before data analysis. | No | An early version of the meta-analysis predictions and preliminary dataset were submitted for a grade in A. E.W's meta-analysis course held at Auburn University. |
| | 3.2 | Describe deviations from the registered aims and methods | NA | |
| | 3.3 | Justify deviations from the registered aims and methods | NA | |
| Eligibility criteria | 4.1 | Report the specific criteria used for including or excluding studies when screening titles and/or abstracts, and full texts, according to the aims of the systematic review (e.g. study design, taxa, data availability) | Yes | |
| | 4.2 | Justify criteria, if necessary (i.e. not obvious from aims and scope) | Yes | |
| **Checklist item** | **Sub-item number** | **Sub-item** | **Reported by authors?** | **Notes** |
| Finding studies | 5.1 | Define the type of search (e.g. comprehensive search, representative sample) | Yes | |
| | 5.2 | State what sources of information were sought (e.g. published and unpublished studies, personal communications) | Yes | |
| | 5.3 | Include, for each database searched, the exact search strings used, with keyword combinations and Boolean operators | Yes | |
| | 5.4 | Provide enough information to repeat the equivalent search (if possible), including the timespan covered (start and end dates) | Yes | |
| Study selection | 6.1 | Describe how studies were selected for inclusion at each stage of the screening process (e.g. use of decision trees, screening software) | Yes | |
| | 6.2 | Report the number of people involved and how they contributed (e.g. independent parallel screening) | Yes | |
| Data collection process | 7.1 | Describe where in the reports data were collected from (e.g. text or figures) | Yes | |

| Checklist item | Sub-item number | Sub-item | Reported by authors? | Notes |
|---|---|---|---|---|
| | 7.2 | Describe how data were collected (e.g. software used to digitize figures, external data sources) | Yes | |
| | 7.3 | Describe moderator variables that were constructed from collected data (e.g. number of generations calculated from years and average generation time) | Yes | |
| | 7.4 | Report how missing or ambiguous information was dealt with during data collection (e.g. authors of original studies were contacted for missing descriptive statistics, and/or effect sizes were calculated from test statistics) | Yes | |
| | 7.5 | Report who collected data | Yes | |
| | 7.6 | State the number of extractions that were checked for accuracy by co-authors | Yes | |

| Checklist item | Sub-item number | Sub-item | Reported by authors? | Notes |
|---|---|---|---|---|
| Data items | 8.1 | Describe the key data sought from each study | Yes | |
| | 8.2 | Describe items that do not appear in the main results, or which could not be extracted due to insufficient information | Yes | |
| | 8.3 | Describe main assumptions or simplifications that were made (e.g. categorising both ‘length’ and ‘mass’ as ‘morphology’) | Yes | |
| | 8.4 | Describe the type of replication unit (e.g. individuals, broods, study sites) | Yes | |
| Assessment of individual study quality | 9.1 | Describe whether the quality of studies included in the systematic review or meta-analysis was assessed (e.g. blinded data collection, reporting quality, experimental *versus* observational) | Yes | Outlier estimates were re-assessed for accuracy by IPG. The three primary models were also re-run without outliers to quantify their influence on overall findings. |
| | 9.2 | Describe how information about study quality was incorporated into analyses (e.g. meta-regression and/or sensitivity analysis) | Yes | |
| Effect size measures | 10.1 | Describe effect size(s) used | Yes | |
| | 10.2 | Provide a reference to the equation of each calculated effect size (e.g. standardised mean difference, log response ratio) and (if applicable) its sampling variance | Yes | |

| | | | |
|---|---|---|---|
| | 10.3 | If no reference exists, derive the equations for each effect size and state the assumed sampling distribution(s) | Yes |
| Missing data | 11.1 | Describe any steps taken to deal with missing data during analysis (e.g. imputation, complete case, subset analysis) | Yes |
| | 11.2 | Justify the decisions made to deal with missing data | Yes |
| Meta-analytic model description | 12.1 | Describe the models used for synthesis of effect sizes | Yes |
| | 12.2 | The most common approach in ecology and evolution will be a random-effects model, often with a hierarchical/multilevel structure. If other types of models are chosen (e.g. common/fixed effects model, unweighted model), provide justification for this choice | Yes |

| **Checklist item** | **Sub-item number** | **Sub-item** | **Reported by authors?** | **Notes** |
|---|---|---|---|---|
| Software | 13.1 | Describe the statistical platform used for inference (e.g. *R*) | Yes | |
| | 13.2 | Describe the packages used to run models | Yes | |
| | 13.3 | Describe the functions used to run models | Yes | |
| | 13.4 | Describe any arguments that differed from the default settings | Yes | |
| | 13.5 | Describe the version numbers of all software used | Yes | |
| Non-independence | 14.1 | Describe the types of non-independence encountered (e.g. phylogenetic, spatial, multiple measurements over time) | Yes | |
| | 14.2 | Describe how non-independence has been handled | Yes | |
| | 14.3 | Justify decisions made | Yes | |
| Meta-regression and model selection | 15.1 | Provide a rationale for the inclusion of moderators (covariates) that were evaluated in meta-regression models | Yes | |
| | 15.2 | Justify the number of parameters estimated in models, in relation to the number of effect sizes and studies (e.g. interaction terms were not included due to insufficient sample sizes) | Yes | |
| | 15.3 | Describe any process of model selection | Yes | |

| | | | | |
|---|---|---|---|---|
| Publication bias and sensitivity analyses | 16.1 | Describe assessments of the risk of bias due to missing results (e.g. publication, time-lag, and taxonomic biases) | Yes | |
| | 16.2 | Describe any steps taken to investigate the effects of such biases (if present) | Yes | |
| | 16.3 | Describe any other analyses of robustness of the results, e.g. due to effect size choice, weighting or analytical model assumptions, inclusion or exclusion of subsets of the data, or the inclusion of alternative moderator variables in meta-regressions | Yes | |
| Clarification of *post hoc* analyses | 17.1 | When hypotheses were formulated after data analysis, this should be acknowledged. | NA | |
| **Checklist item** | **Sub-item number** | **Sub-item** | **Reported by authors?** | **Notes** |
| Metadata, data, and code | 18.1 | Share metadata (i.e. data descriptions) | Yes | |
| | 18.2 | Share data required to reproduce the results presented in the manuscript | Yes | |
| | 18.3 | Share additional data, including information that was not presented in the manuscript (e.g. raw data used to calculate effect sizes, descriptions of where data were located in papers) | Yes | |
| | 18.4 | Share analysis scripts (or, if a software package with graphical user interface (GUI) was used, then describe full model specification and fully specify choices) | Yes | |
| Results of study selection process | 19.1 | Report the number of studies screened | Yes | |
| | 19.2 | Report the number of studies excluded at each stage of screening | Yes | |
| | 19.3 | Report brief reasons for exclusion from the full text stage | Yes | |
| | 19.4 | Present a Preferred Reporting Items for Systematic Reviews and Meta-Analyses (PRISMA)-like flowchart (www.prisma-statement.org). | Yes | |

| | | | | |
|---|---|---|---|---|
| Sample sizes and study characteristics | 20.1 | Report the number of studies and effect sizes for data included in meta-analyses | Yes | |
| | 20.2 | Report the number of studies and effect sizes for subsets of data included in meta-regressions | Yes | |
| | 20.3 | Provide a summary of key characteristics for reported outcomes (either in text or figures; e.g. one quarter of effect sizes reported for vertebrates and the rest invertebrates) | Yes | |
| | 20.4 | Provide a summary of limitations of included moderators (e.g. collinearity and overlap between moderators) | Yes | |
| | 20.5 | Provide a summary of characteristics related to individual study quality (risk of bias) | Yes | |
| **Checklist item** | **Sub-item number** | **Sub-item** | **Reported by authors?** | **Notes** |
| Meta-analysis | 21.1 | Provide a quantitative synthesis of results across studies, including estimates for the mean effect size, with confidence/credible intervals | Yes | |
| Heterogeneity | 22.1 | Report indicators of heterogeneity in the estimated effect (e.g. $I^2$, $tau^2$ and other variance components) | Yes | |
| Meta-regression | 23.1 | Provide estimates of meta-regression slopes (i.e. regression coefficients) and confidence/credible intervals | Yes | |
| | 23.2 | Include estimates and confidence/credible intervals for all moderator variables that were assessed (i.e. complete reporting) | Yes | |
| | 23.3 | Report interactions, if they were included | Yes | |
| | 23.4 | Describe outcomes from model selection, if done (e.g. R2 and AIC) | NA | |
| Outcomes of publication bias and sensitivity analyses | 24.1 | Provide results for the assessments of the risks of bias (e.g. Egger's regression, funnel plots) | Yes | |
| | 24.2 | Provide results for the robustness of the review's results (e.g. subgroup analyses, meta-regression of study quality, results from alternative methods of analysis, and temporal trends) | Yes | |

| | | | | |
|---|---|---|---|---|
| Discussion | 25.1 | Summarise the main findings in terms of the magnitude of effect | Yes | |
| | 25.2 | Summarise the main findings in terms of the precision of effects (e.g. size of confidence intervals, statistical significance) | Yes | |
| | 25.3 | Summarise the main findings in terms of their heterogeneity | Yes | |
| | 25.4 | Summarise the main findings in terms of their biological/practical relevance | Yes | |
| | 25.5 | Compare results with previous reviews on the topic, if available | Yes | |
| | 25.6 | Consider limitations and their influence on the generality of conclusions, such as gaps in the available evidence (e.g. taxonomic and geographical research biases) | Yes | |
| **Checklist item** | **Sub-item number** | **Sub-item** | **Reported by authors?** | **Notes** |
| Contributions and funding | 26.1 | Provide names, affiliations, and funding sources of all co-authors | Yes | |
| | 26.2 | List the contributions of each co-author | Yes | |
| | 26.3 | Provide contact details for the corresponding author | Yes | |
| | 26.4 | Disclose any conflicts of interest | NA | No COIs to report. |
| References | 27.1 | Provide a reference list of all studies included in the systematic review or meta-analysis | Yes | |
| | 27.2 | List included studies as referenced sources (e.g. rather than listing them in a table or supplement) | No | Sourced studies omitted from official reference list on account of space limitations. |
