## Appendix S3 for "The fitness consequences of wildlife conservation translocations: a meta-analysis"

to

**CONTENTS**

**Supplementary Figure S1: Time-lag bias with outliers excluded** ...................................... **p. 2**

**Supplementary Figure S2: Egger's regression with outliers excluded** ............................. **p. 3**

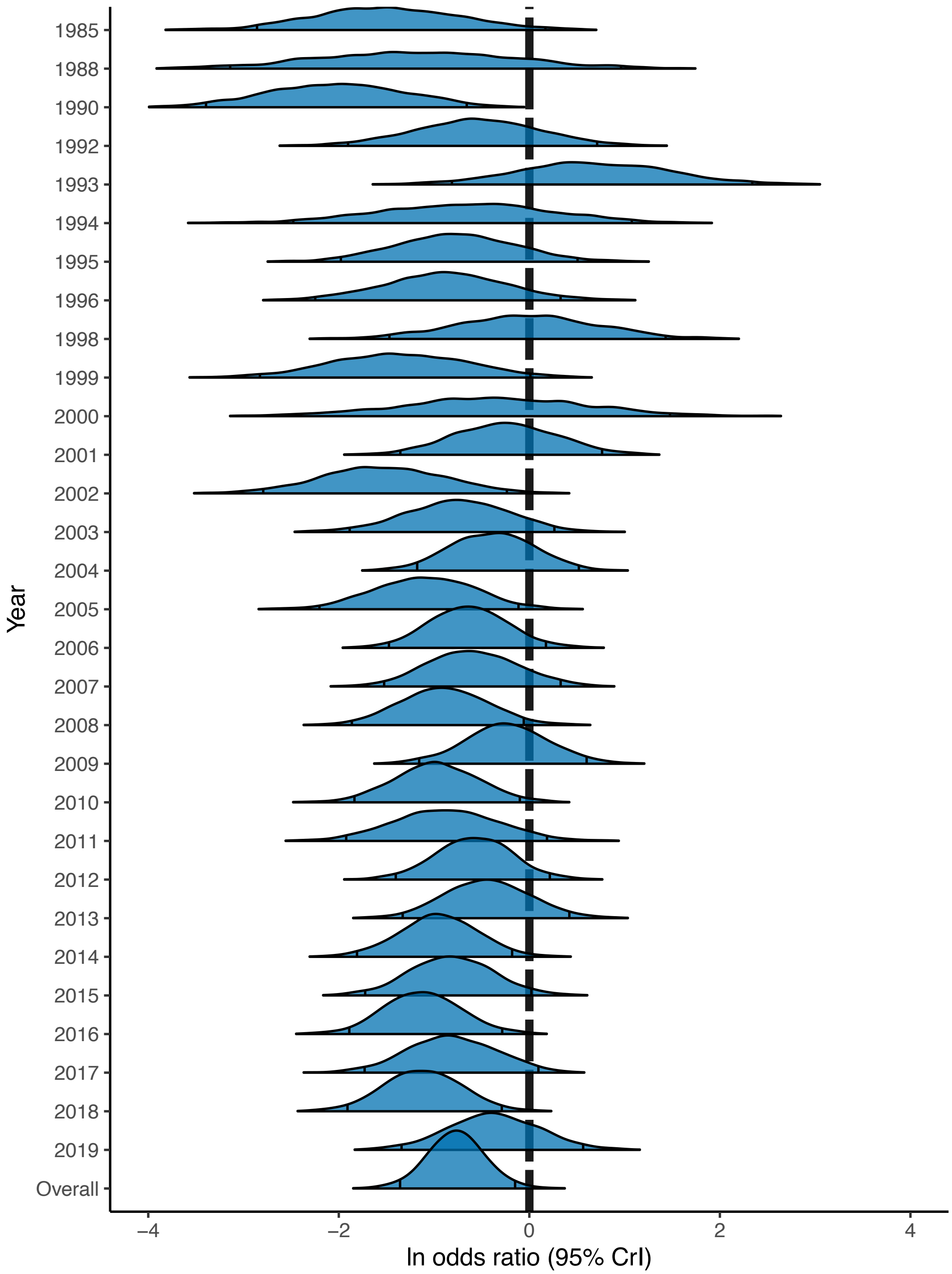


**Supplementary Figure 1.** Time-lag bias. The relationship between effect size and publication year, with outliers excluded. Effect sizes are marginal posterior distributions, with 95% credible intervals indicated by black vertical lines. The dashed vertical line indicates an effect size of zero.

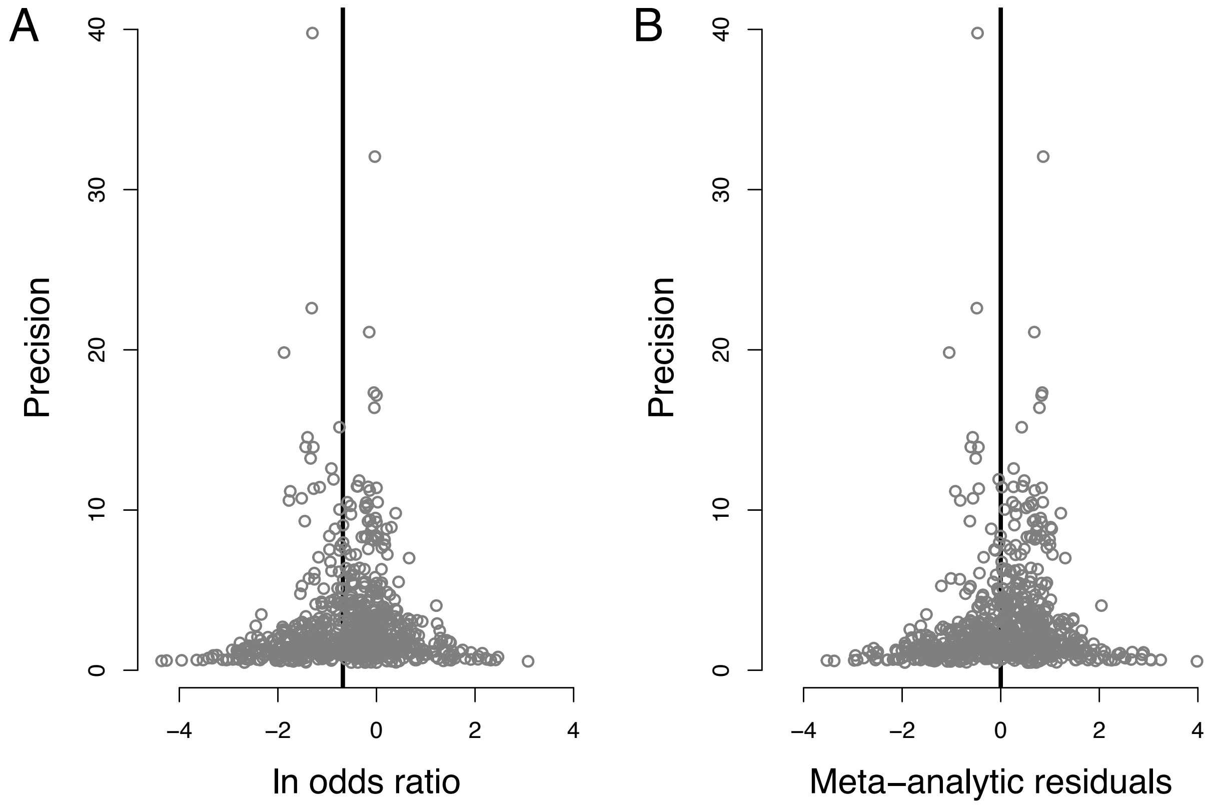


**Supplementary Figure 2.** Egger's regression. Funnel plots describing patterns of publication bias for the full dataset excluding outliers. (A) Relationship between effect size (log odds ratio) and precision. The solid vertical line indicates the meta-analytic mean. (B) Relationship between the meta-analytic residuals and precision. The solid vertical line indicates zero.
